# Dynamic brain-to-brain concordance and behavioral mirroring as a mechanism of the patient-clinician interaction

**DOI:** 10.1101/2020.08.05.237511

**Authors:** Dan-Mikael Ellingsen, Kylie Isenburg, Changjin Jung, Jeungchan Lee, Jessica Gerber, Ishtiaq Mawla, Roberta Sclocco, Karin B Jensen, Robert Randolph Edwards, John M. Kelley, Irving Kirsch, Ted J. Kaptchuk, Vitaly Napadow

## Abstract

The patient-clinician interaction can powerfully shape treatment outcomes such as pain, but is often considered an intangible “art-of-medicine”, and has largely eluded scientific inquiry. Although brain correlates of social processes such as empathy and theory-of-mind have been studied using single-subject designs, the specific behavioral and neural mechanisms underpinning the patient-clinician interaction are unknown. Using a two-person interactive design, we simultaneously recorded functional MRI (i.e. hyperscanning) in patient-clinician dyads, who interacted via live video while clinicians treated evoked pain in chronic pain patients. Our results show that patient analgesia is mediated by patient-clinician nonverbal behavioral mirroring and brain-to-brain concordance in circuitry implicated in theory-of-mind and social mirroring. Dyad-based analyses showed extensive dynamic coupling of these brain nodes with the partners’ brain activity, yet only in dyads where clinical rapport had been established prior to the interaction. These findings point to a putatively key brain-behavioral mechanism for therapeutic alliance and psychosocial analgesia.

---

The patient-clinician interaction is fundamental to clinical care. Positive clinical encounters are associated with higher patient satisfaction, mutual trust (*1*), treatment adherence (*2*), and even clinical outcomes (*3*–*5*). Conversely, suboptimal interactions may propagate miscommunication (*6*), clinician burnout (*7*), patient distrust (*8*), and discourage care seeking (*9*). The patient-clinician relationship is also likely to account for a substantial part of psychologically mediated relief (e.g. placebo analgesia) (*10*). Yet, clinical engagement is often considered an intangible “art-of-medicine,” and scientific inquiry into the specific underpinning mechanisms has been minimal. A scientific understanding of the neurobiological and behavioral mechanisms supporting the patient-clinician interaction may be key to harnessing this untapped potential to improve clinical care.

A number of neuroimaging studies have established that brain regions including TemporoParietal Junction, anterior insula, and ventrolateral Prefrontal Cortices are implicated in social processes such as empathy and theory-of-mind (inferring the mental state of others) (*11*), which may also be relevant for the clinical encounter. Indeed, a recent study indicated that this brain circuitry is activated in clinicians applying pain treatment to individuals appearing to be in pain (*12*). However, while most neuroimaging studies have employed single-subject experimental designs, it is increasingly recognized that understanding the complex neural dynamics of social interactions, such as in the clinical dyad, requires the investigation of simultaneous brain activity in patients and clinicians during actual interaction (*13*).

For example, a large literature points to behavioral mirroring and physiological concordance as fundamental to human affiliation and bonding (*14*, *15*). In the context of clinical interaction, verbal (*16*) and nonverbal (*17*) behavioral synchrony between patients and clinicians is associated with better therapeutic effectiveness and relationship quality (*18*). Furthermore, concordance in sympathetic nervous system activation has been associated with higher physician empathy and less emotional distance (*19*).

Recent functional brain imaging studies of two (or more) people during interaction (i.e. hyperscanning) have found that activity in social mirror networks synchronizes between individuals when socially interacting (*20*, *21*), and stronger coupling may reflect more successful communication (*22*), suggesting that concordance of brain activity in social mirroring networks may play a key role in social interaction (*13*, *23*).

Here, we investigated patient-clinician mirroring in facial expressions and dynamic brain activity concordance as potential mechanisms supporting clinical outcomes mediated by patient-clinician interactions. We used functional Magnetic Resonance Imaging (fMRI) to record brain activity simultaneously (fMRI hyperscanning) in chronic pain patients and clinicians (acupuncturists) during an ecologically-valid yet experimentally-controlled clinical encounter, in which the clinician treated the patient to reduce evoked pain (**Fig. 1a**).

**Fig. 1:**
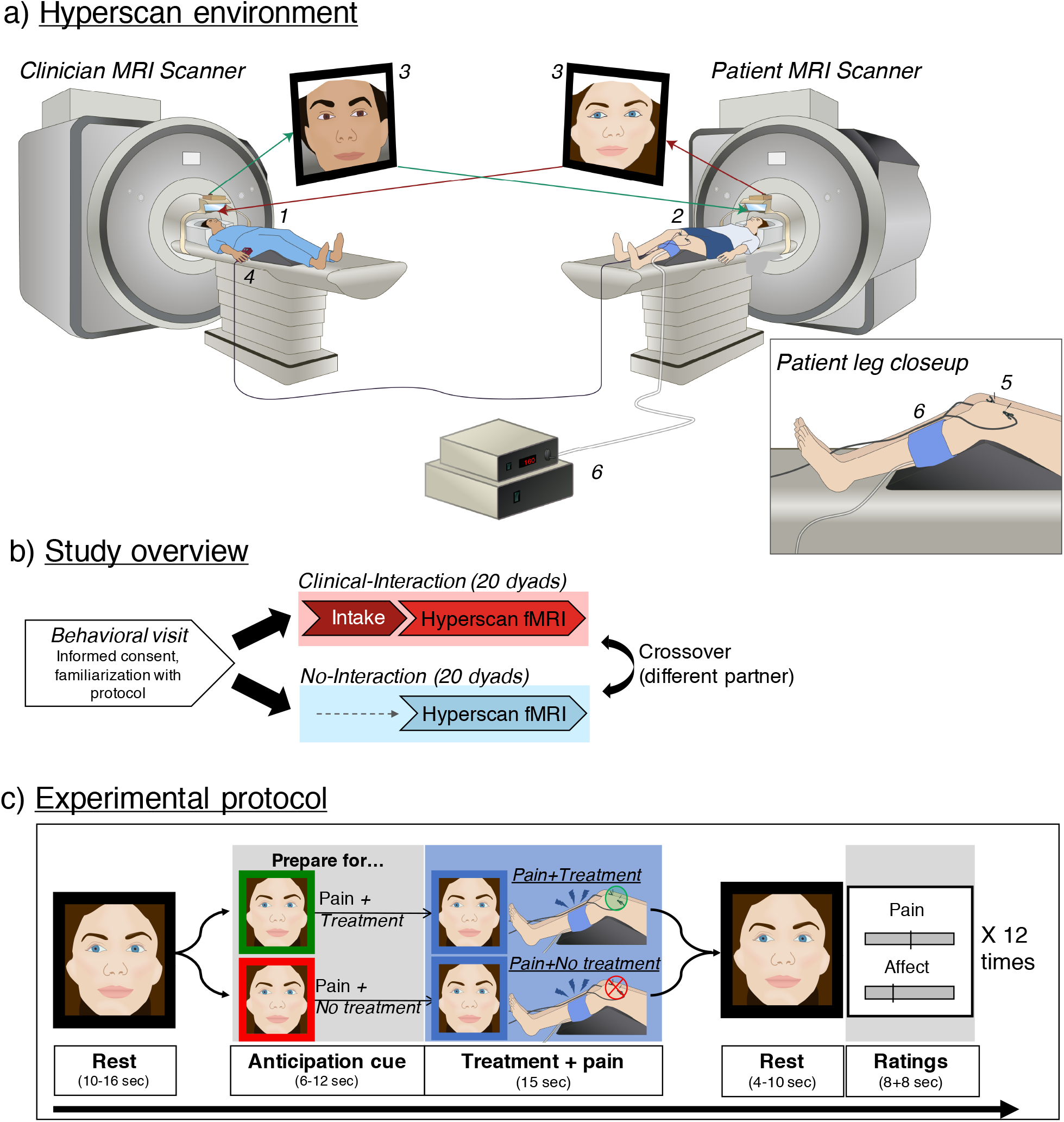
Study setup. a) The fMRI hyperscanning environment. The clinician (1) and patient (2) were positioned in two different 3T MRI scanners. An audio-video link enabled on-line communication between the two scanners (3), and video images were used to frame-by-frame facial expression metrics. During simultaneous acquisition of BOLD-fMRI data, the clinician used a button box (4) to apply electroacupuncture treatment (real/sham, double-blind) to the patient (5) to alleviate evoked pressure pain to the leg (6, Hokanson cuff inflation). Pain and affect related to the treatment were rated after each trial. b) Study overview. After an initial behavioral visit, each subject participated in a Clinical-Interaction (Hyperscan preceded by a clinical intake) and No-Interaction condition (Hyperscan without a preceding intake), in a counterbalanced order, with two different partners. c) Experimental protocol. Each hyperscan was composed of 12 repeated trials (4 verum Electroacupuncture (EA), 4 sham EA, 4 no treatment) in a pseudorandomized order. After a resting period (far left), both participants were shown a visual cue to indicate whether the next pain stimulus would be treated (green frame) ornot treated (red frame) by the clinician. These cues prompted clinicians prepare to either apply ornotapply treatment, while evoking corresponding anticipation for the patient. Following the anticipation cue, moderately painful pressure pain was applied to the patient’s left leg, while the clinician applied or did not apply treatment, respectively. After another resting period, participants rated pain (patients), vicarious pain (clinicians), and affect (both) using a Visual Analog Scale.

We enrolled 45 participants, including 23 female chronic pain patients diagnosed with fibromyalgia for at least one year and 22 acupuncture clinicians (15 female). Each participant was matched with up to 2 partners, forming a total of 40 distinct, interacting dyads. Each dyad was scanned under one of two conditions (counterbalanced order): in the ‘Clinical-Interaction’ condition, the clinician performed a clinical consultation and intake with the patient prior to MRI scanning to enable the dyad to establish clinical rapport. The ‘No-Interaction’ control condition was identical to ‘Clinical-Interaction’ except that the patient and the clinician had not had an intake, and were only introduced briefly before scanning (**Fig. 1b**). Due to data loss, we obtained complete MRI data from 37 dyads. See Methods for comprehensive methodological details.

## Results

### Therapeutic alliance

Each participant completed 4 sessions: 1) A behavioral session for informed consent and familiarization with protocol; 2) a clinical intake session, in which the clinician (acupuncturist) performed an intake with the patient, encouraged to be ‘as similar as possible to your daily practice’ to maximize ecological validity; 3) a Clinical-Interaction MRI on a separate day after the intake, in which the same patient and clinician were scanned together during a pain treatment session; and 4) a No-Interaction MRI to control for the social relationship established at the intake. The order of Clinical-Interaction MRI and No-Interaction MRI was counter-balanced between subjects (**Fig. 1b**). Different MRI visits were always on separate days.

The Consultation And Relational Empathy (CARE) (24) scale was collected after each session as a proxy for therapeutic alliance. A repeated-measures ANOVA confirmed that patients reported different levels of therapeutic alliance depending on the context of the dyadic clinical in-teraction (F(1.34,18.76)=20.82, P<0.001, η_p_^2^=0.60). Planned direct comparisons indicated significantly lower CARE scores for No-Interaction MRI (Mean±SD=32.19±8.09), compared to Intake (42.20±4.25, t=5.84, P<0.001, Cohen’s *d*=1.46, 95% confidence interval [CI]=5.39,14.61) and Clinical-Interaction MRI (41.63±5.11, t=5.21, P<0.001, d=1.30, CI=4.56,14.32) con-texts (**Fig. S1**). No significant difference was noted between Intake and Clinical-Interaction MRI (t=0.63, P=0.54, d=0.16, CI=−1.85,2.97) sessions. A similar pattern was seen for clinician-rated therapeutic alliance (**see Fig. S1**).

### Evoked pressure pain, vicarious pain, and treat-ment-related affect

MRI-compatible video cameras allowed participants to communicate non-verbally (e.g. eye movement, facial expressions) throughout hyperscanning. During block-design fMRI, patients received 12 moderately painful cuff pressures to the left leg (**Fig. 1c,** see Supplementary Methods for details on stimulus presentation). Importantly, enrolling acupuncture practitioners as clinicians allowed for therapy to be administered *during* hyperscanning, using remote, but ecologically-valid, controlled electroacupuncture (EA) through two needles placed above the patients’ knee (pseudorandomized verum, sham, and overt No-Treatment, 15 s duration). Prior to each pain stimulus, both participants were given a visual cue (6-12 s jittered, frame around face changing color) indicating whether upcoming pain stimuli would be accompanied by Treatment (green) or No-Treatment (red). For patients, this cue elicited an anticipation of receiving or not receiving treatment for the upcoming pain, whereas for clinicians this prompted them to prepare for whether-or-not to apply treatment. During cuff inflation, the clinician correspondingly pressed and held either the ‘Treatment’ button or a different ‘No treatment control’ button. After each stimulus (4-10 s jittered), the patients and clinicians rated pain intensity (patients), vicarious pain (clinicians), and affect (patients and clinicians) using Visual Analog Scales.

There was no significant difference in pain between sham and verum EA (t=0.83, P=0.42. Therefore, these conditions were pooled together as ‘Treatment’, collectively, for further analyses, and Treatment – No-Treatment differences are referred to as ‘analgesia’ (see Methods). For patients’ pain intensity, a repeated measures ANOVA confirmed a main effect of ‘Treatment condition’, in which pain intensity was rated significantly lower for Treatment (Mean±SD=26.32±15.92), relative to No-Treatment (32.94±17.98, F(1,15)=9.79, P=0.007, η_p_^2^=0.40, CI=1.02,12.22, **Fig. 2a**). There was no main effect of ‘Clinical context’ (levels: Clinical-Interaction and No-Interaction, F(1,15)=0.04, P=0.84, η_p_^2^=0.003, CI=−8.03,8.03) and no statistical interaction between ‘Clinical context’ and ‘Treatment condition’ (F(1,15)<0.01, P=0.98, η_p_^2^<0.01), suggesting pain intensity and analgesia were comparable across different clinical interaction contexts. Further, there were no interactions involving ‘Order’ (P’s>0.12). However, an ANCOVA confirmed that patient analgesia was significantly associated with subjective evaluations of the relationship, indicating that in dyads where relationship quality was rated more highly, patients reported stronger analgesia (‘HRS score’, F(1,295)=7.36, P=0.007, η_p_^2^=0.02). There were no significant main effects or interactions involving clinical interaction context (P’s>0.06) and ‘HRS item’ (P’s>0.72), suggesting the association between analgesia and relationship evaluation was comparable across HRS items and clinical interaction contexts.

**Fig. 2:**
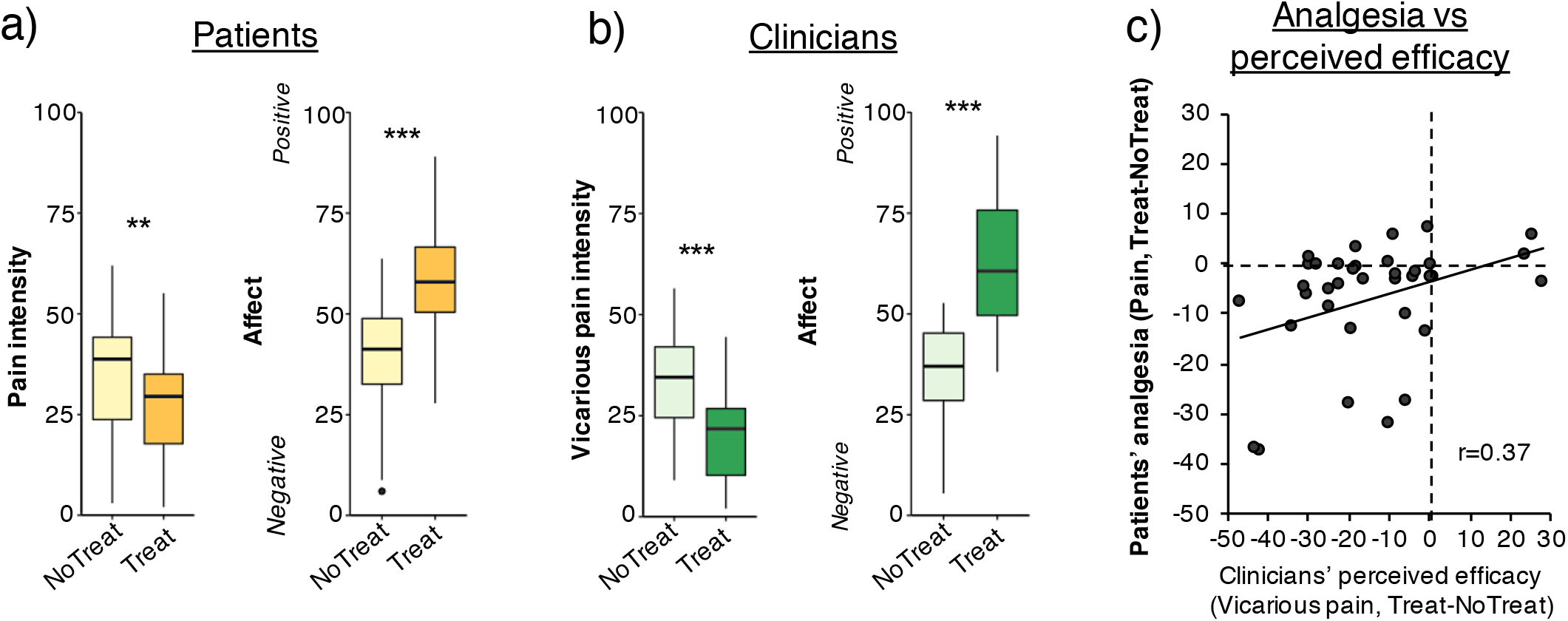
Self-reported pain and affect during fMRI hyperscanning. A) Patients reported less pressure pain intensity when being treated by the clinician, relative to no-treatment trials. Furthermore, they reported feeling more positive affect while treated, relative to no-treatment trials (VAS rating, «How did you feel about (not) getting treated with electroacupuncture?», anchors: Extremely negative/positive). B) Correspondingly, clinicians thought patients had less pain during treatment trials relative to no-treatment trials, and they reported more positive affect while treating relative to not treating (VAS rating, «How did you feel about (not) doing the electro-acupuncture?», anchors: Extremely positive/negative. C) A correlation between patients’ ΔPain (Treat-NoTreat) and clinicians’ ΔVicarious pain (Treat-NoTreat) difference scores suggested that for patients who reported greater pain relief, their clinician also perceived higher treatment efficacy. VAS: Visual Analog Scale. **p<0.01; ***p<0.005

For clinicians’ ratings of vicarious pain there was a main effect of ‘Treatment condition’, in which vicarious pain was rated as significantly lower for Treatment (18.52±13.62) relative to No-Treatment (33.06±18.79, F(1,15)=17.27, P<0.001, η_p_^2^=0.55, CI=7.27,21.82, **Fig. 2b**). There was no main effect of ‘Clinical context’ (F(1,15)=0.51, P=0.49, η_p_^2^=0.04, CI=−7.19,14.70), no statistical interaction with treatment condition (F(1,15)=0.51, P=0.47, η_p_^2^ =0.04) and no interactions involving ‘Order’ (P’s>0.61). Furthermore, patients’ analgesia (ΔPain, Treat-NoTreat) correlated with clinicians’ perceived treatment efficacy (ΔVicarious pain, Treat-No-Treat), such that for patients who reported greater pain relief, their clinician also perceived higher treatment efficacy (r=0.37, P=0.02), supporting patients’ ability to communicate their subjective pain to their clinician (**Fig. 2c**).

Correspondingly, repeated-measures ANOVAs on ratings of affect indicated that both patients (F(1,15)=10.69, P=0.005, η_p_^2^=0.416, CI=8.37,29.81) and clinicians (F(1,15)=12.35, P=0.003, η_p_^2^ =0.47, CI=13.18,39.78) felt more positively about Treatment trials than No-Treatment trials (**Fig. 2a-b**), while ratings were comparable across Clinical-Interaction and No-Interaction contexts (Patients: F(1,15)=0.02, P=0.90, η_p_^2^=0.001, CI=-5.25,2.11; clini-cians: F(1,15)=0.01, P=0.92, η_p_^2^ <0.01, CI=-6.48,5.99). There were no ‘Clinical Context’*’Treatment’ statistical interactions (Patients: F(1,15)=0.57, P=0.46, η_p_^2^=0.001; clinicians: F(1,15)=1.3, P=0.27, η_p_^2^ =0.09), indicating that affect was comparable across scans. There were no significant statistical interactions involving order (Patients: P’s>0.26; Clinicians: P’s>0.11).

### Facial mirroring was associated with placebo analgesia and therapeutic alliance

In-scanner videos were recorded and processed using automated facial feature (expression) extraction (Affectiva, Cambridge, MA). Average values for individual features were calculated for each trial. To assess treatment-related change in facial mirroring, we then calculated the correlation coefficient (r-to-z transformed) between patients and clinicians for the Treat – No-treat change score across all features, resulting in one overall facial mirroring score per dyad (**Fig. 3a**). During anticipation of pain, facial mirroring across expressions correlated significantly with therapeutic alliance at MRI (r=0.51, P=0.036) and patients’ ratings of analgesia (r=-0.52, P=0.031, **Fig. 3b**).

**Fig. 3:**
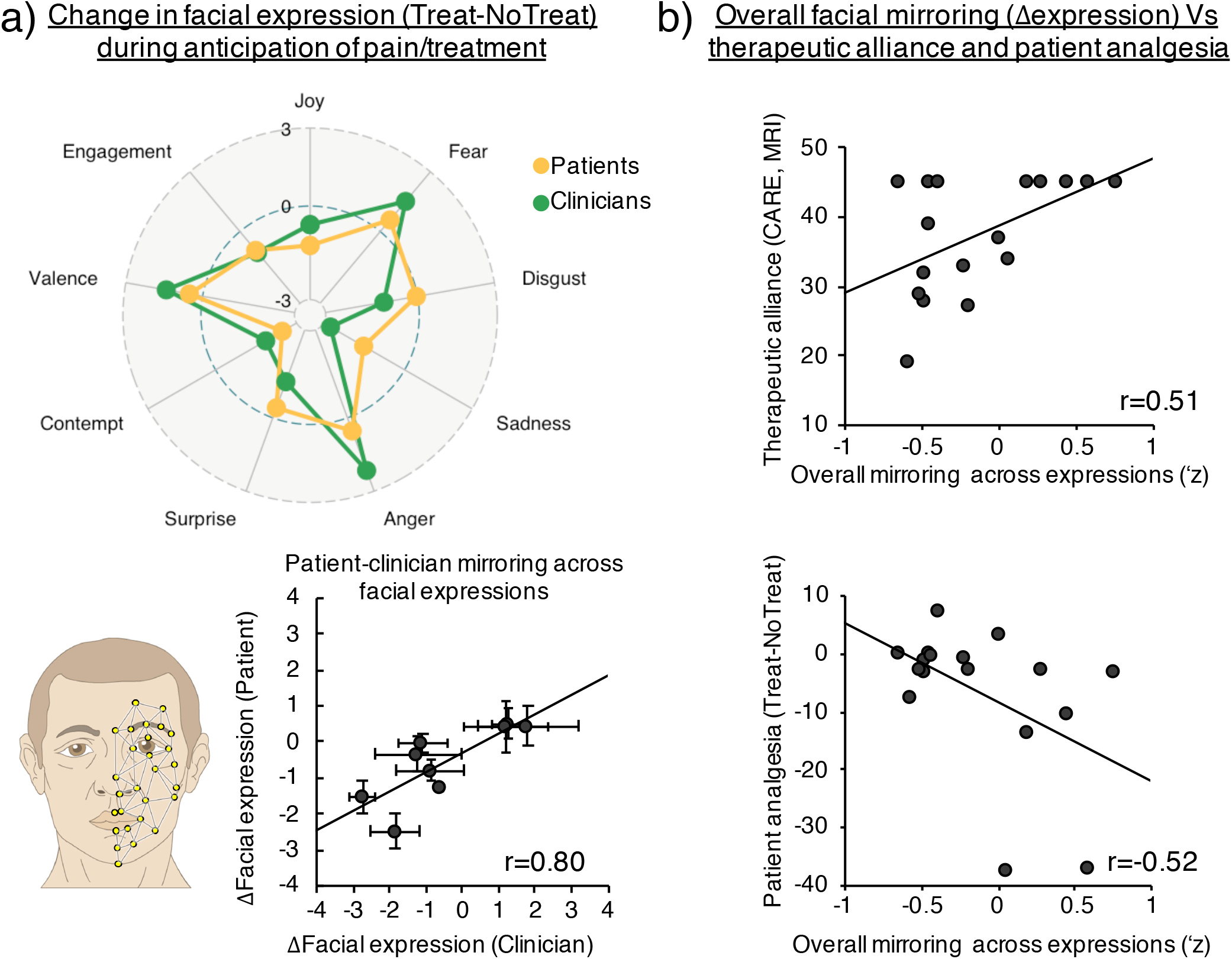
Patient-clinician mirroring in facial expressions during the therapeutic encounter. We used automated detection of facial muscle units, which were used to calculate frame-by-frame emotional expression scores (Affectiva, Cambridge, MA). a) For the whole group, we found strong correspondence in treatment-induced change (Treat-NoTreat) between patients and clinicians across expressions. b) To assess facial mirroring across expressions, we calculated the Treatment-induced (Treat-NoTreat) change for each expression for the patient and clinician, and subsequently Pearson’s coefficients (r-to-z transformed), across expressions within each dyad. This approach enabled higher sensitivity to differences in patterns of expressions between different dyads, compared to e.g. assessing similarity within single expression metrics. We found that increased facial mirroring (overall, across all expressions) was associated with higher therapeutic alliance and stronger patient-reported analgesia (more negative values mean stronger pain reduction).

### Brain activation associated with pain and analgesia

To investigate treatment-related change in brain processing of evoked pressure pain, we first performed a whole-brain group GLM for all patients, for the contrast ‘Treatment’ – ‘No-Treatment’, which indicated increased fMRI activation of bilateral vlPFC, TPJ, dlPFC, and mPFC, in addition to left STS for treated, relative to nontreated, pain (**Fig. S2**). We then investigated brain circuitry associated with individual treatment analgesia in the patients’ brain. A whole-brain regression analysis showed that stronger analgesia (NoTreat-Treat pain ratings) was associated with greater treatment-related increase in patients’ right vlPFC, precuneus, visual circuitry, and a cluster in the Inferior Parietal Lobule (IPL)/Supramarginal Gyrus (SMG) during pain (Treat-NoTreat) (**Fig. S3**).social mirroring circuitry regions (e.g. vlPFC) for the patient during pain.

### Shared activation between patients and clinicians in brain circuitry associated with social mirroring

Next, we investigated dynamic brain activity concordance between patients and clinicians, focusing on the anticipation period, when the relationship may impact brain activity without competing neural processing of nociceptive afference (for patients) or motor activity for treatment delivery (for clinicians), as during the pain/treatment period.

To assess brain activity concordance, we first calculated brain response to anticipation of pain, collapsed over Treat/NoTreat conditions (**Fig. 4a, left**), as concordance related to therapeutic alliance and pain outcomes could be driven by social interaction during the anticipation of both treated and non-treated pain. Next, we performed a whole-brain voxelwise conjunction analysis using the minimum statistic to investigate brain circuitry commonly activated for both patients and clinicians, which provided regions of interest (ROI) for dynamic concordance analyses. This group conjunction analysis demonstrated shared anticipatory activations between patients and clinicians in bilateral circuitry implicated in social mirroring, theory-of-mind, and social cognition (e.g. bilateral Temporoparietal Junction, TPJ, left ventrolateral Prefrontal Cortex, vlPFC, and left anterior insula, aINS. See **Fig. S4** for analyses of patients’ and clinicians’ brain responses during the pain phase.

**Fig. 4:**
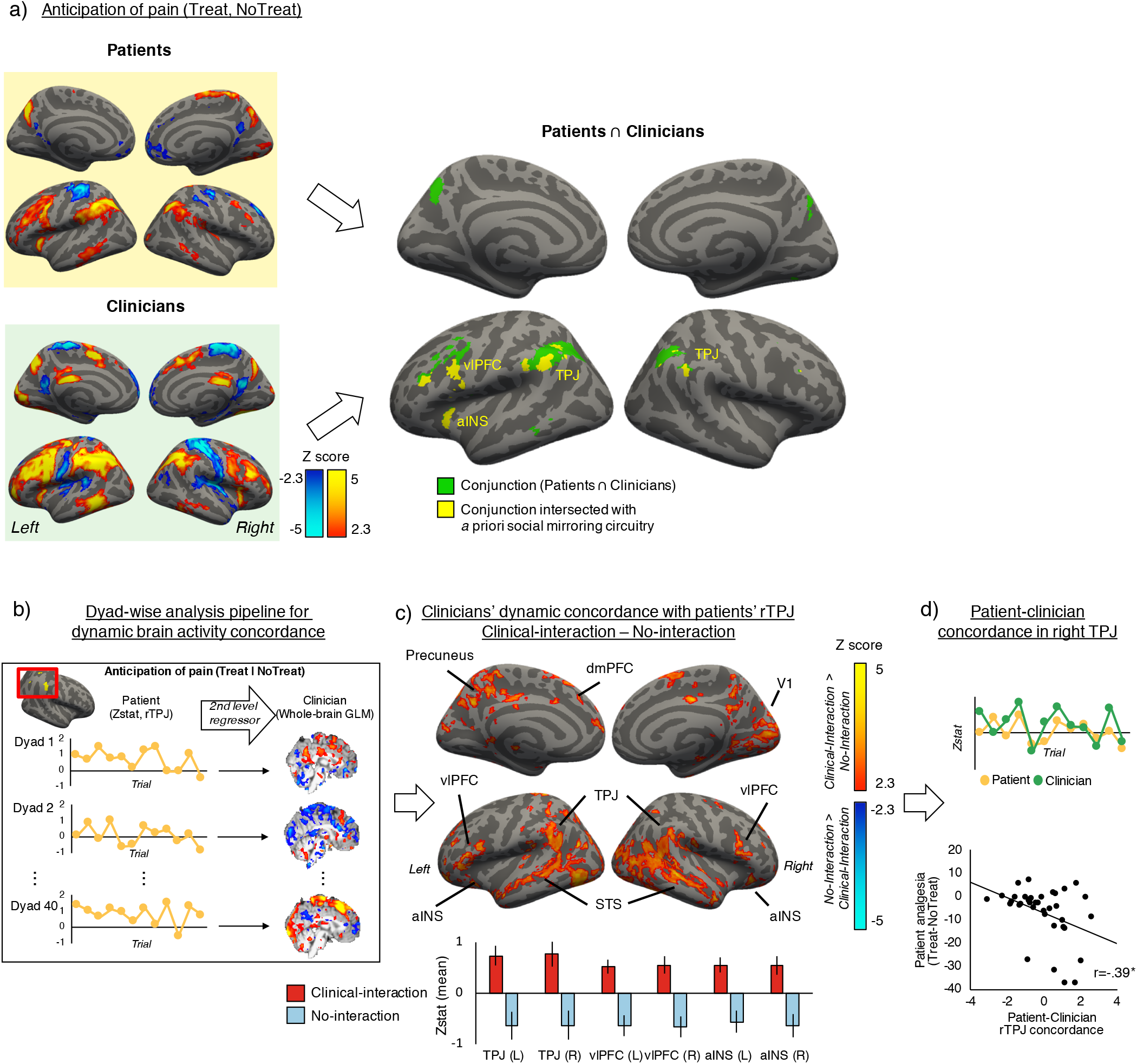
Shared brain activation and dynamic concordance between patients and clinicians. a) The left panel shows fMRI responses to anticipation of receiving pain (patients, top left) and preparing to provide / not provide treatment (clinicians, bottom left). Aconjunction analysis of these activation maps demonstrated common anticipatory activation for patients and clinicians in brain circuitry implicated in social mirroring, theory-of-mind, and empathy, such as left vlPFC, aINS, bilateral TPJ, and left Superior Temporal Sulcus, in addition to the precuneus, a cluster comprising bilateral supramarginal/angular gyrus, and superior parietal lobule. b) To assess dynamic concordance in brain activity between patients and clinicians, throughout the pain/treatment scan, we extracted trial-by-trial Z-scores from the patient’s rTPJ, which were then used as regressors in the clinician’s second-level GLM. This analysis used trial-by-trial whole-brain contrast parameter estimates for the pain/treatment anticipation block. c) A group-level analysis of clinician dynamic concordance with patients’ rTPJ showed that Clinical-Interaction, relative to No-Interaction control, enhanced rTPJ concordance to circuitry implicated in mentalizing, empathy, and social mirroring, e.g. TPJ, vlPFC, aINS, and STS, in addition to visual circuitry and precuneus, a key node of the default mode network. The bottom panel shows mean Z-statistical values from extracted ROIs, with error bars indicating SEM. d) For the enhanced rTPJ-to-rTPJ contrast, dynamic fMRI response was indeed driven by increased concordance between patients and clinicians following Clinical-Interaction (top, example dyad). Greater patient-clinician rTPJ concordance was associated with stronger patient analgesia (bottom). rTPJ=right Temporoparietal Junction, vlPFC=ventrolateral Prefrontal Cortex, aINS=anterior Insula, STS=Superior Temporal Sulcus, fMRI=functional Magnetic Resonance Imaging.

### Social interaction enhanced patient-clinician dynamic concordance in brain activity

For each dyad, we then extracted each individual’s mean ROI Z-statistical value from each trial, which were used as a regressor in a second-level GLM for their dyadic partner’s fMRI data, providing a whole-brain map of dynamic concordance with the partner’s ROIs for each dyad (**See Methods and Fig. S5 for details**). Employing a dynamic metric is important as concordance is best defined by shared deviations in brain response across dyad members (*25*). Following Clinical-Interaction, dynamic (trial-to-trial) rTPJ_Patients_ concordance was evident with clinicians’ brain response in circuitry implicated in social mirroring, theory-of-mind, and social cognition (e.g. bilateral TPJ, vlPFC, aINS), in addition to visual and executive control circuitry, and significantly differed from the No-Interaction context for these regions (**Fig. 4b-c**). ROI extraction from the clinicians’ whole-brain maps demonstrated that concordance between patients’ and clinicians’ rTPJ (but not other ROIs from above, r’s=−0.11–0.17, P’s>0.5) was significantly associated with patients’ analgesia (r=-0.39, P=0.017, **Fig. 4d**). Analyses exploring effects of Clinical-Interaction on dynamic concordance with other nodes of the social mirroring circuitry are shown in **Fig. S6)**.

### Patients’ treatment-related brain response to pain mediated the effect of rTPJ concordance on analgesia

Finally, we explored whether concordance effects on analgesia were mediated by treatment-related change in specific social mirroring circuitry regions (e.g. vlPFC) for the patient *during* pain. We found that stronger treatment analgesia was associated with increased treatment-induced fMRI response in pain modulatory circuitry, e.g. vlPFC (**Fig. S3**). The bootstrapped mediation analysis indicated a significant effect of the indirect path (a*b=-1.80, P=0.006, CI=-3.90,-0.47), indicating that treatment-related change in patients’ vlPFC response during pain (Pain_Treat-NoTreat_) mediated the effect of anticipatory patient-clinician rTPJ concordance on analgesia (**Fig. 5**). Other nodes in the social mirroring circuitry activated during Pain_Treat-_ NoTreat did not significantly mediate this relationship (P’s>0.07).

**Fig. 5:**
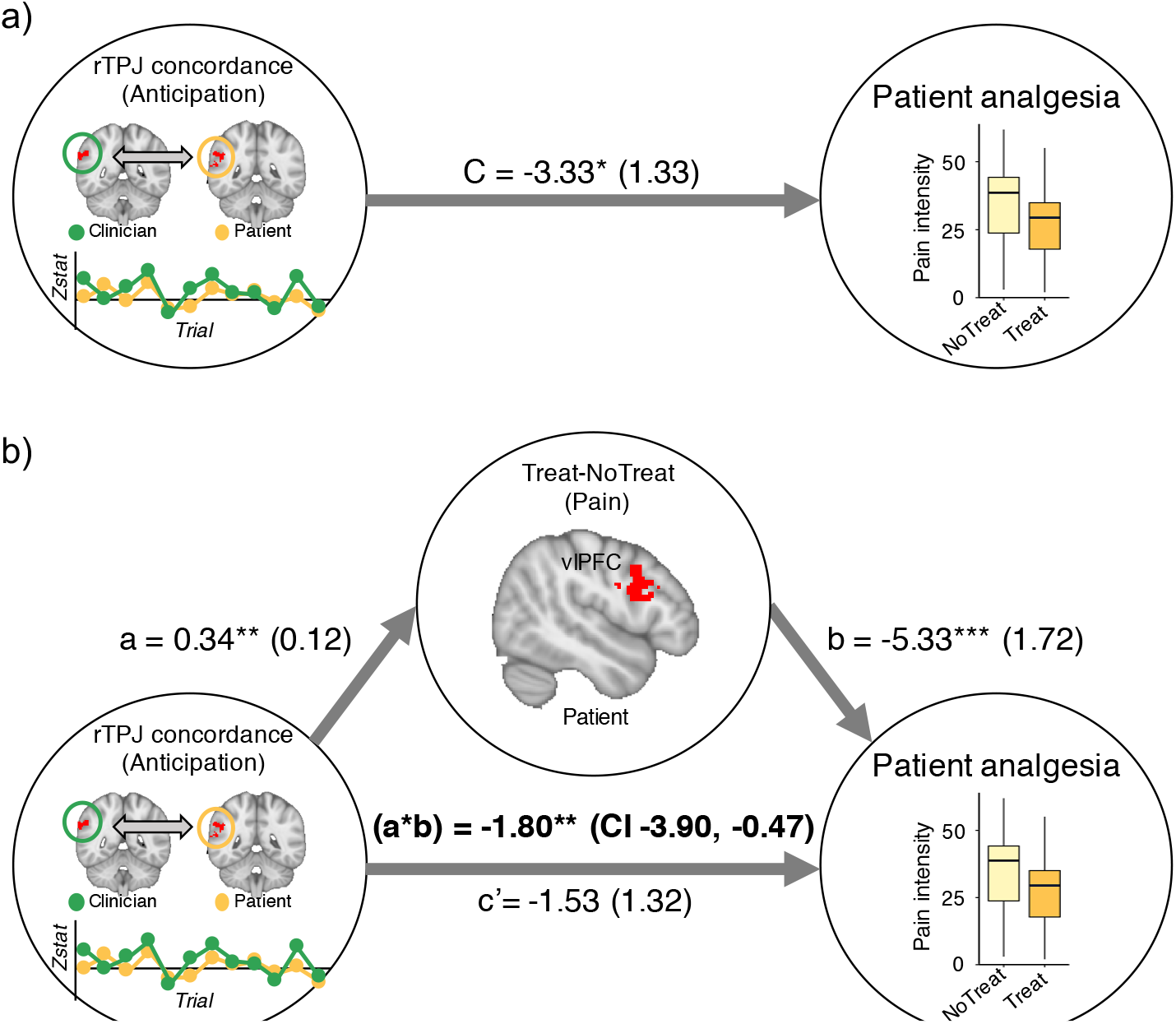
Patients’ treatment-related change in vlPFC mediated theassociation between rTPJ concordance and analgesia. A) Anticipatory rTPJ concordance between patients and clinicians showed a direct linear association with patient analgesia. B) A mediation analysis showed that Treatment-related change in patients’ vlPFC response during pressure pain statistically mediated the association between rTPJ concordance and patient analgesia, suggesting a mechanism in which patient-clinician rTPJ concordance recruits a pain modulatory vlPFC response in the patient’s brain. rTPJ=right Temporoparietal Junction; vlPFC = ventrolateral Prefrontal Cortex

## Discussion

We identified a putative brain-behavioral mechanism supporting the patient-clinician relationship and how it may influence clinical outcomes. We found that dynamic patient-clinician concordance in brain activity implicated in social mirroring and theory-of-mind was increased after the establishment of therapeutic alliance through a clinical interaction. Furthermore, stronger brain concordance was associated with stronger analgesia, an association that was mediated by activation of pain modulatory circuitry in the patient during pain. Finally, increased facial mirroring between patients and clinicians was associated with stronger therapeutic alliance and greater analgesia.

Patient-clinician behavioral synchrony and reciprocity is thought to support processes such as mutual empathy and therapeutic alliance (*26*), and thus constitutes a cornerstone for patient-centered care (*18*, *27*). We found that circuitry implicated in social mirroring (TPJ, vlPFC, and aINS) was commonly activated in both patients and clinicians during anticipation of pain and treatment. Dyad-based analyses suggested that these nodes showed extensive dynamic coupling with the partners’ brain activity, but only in dyads who had established a clinical relationship prior to MRI. Specifically, rTPJ-to-rTPJ concordance showed the strongest association with patients’ analgesia. The TPJ is a key hub in theory-of-mind processes, i.e. mentalizing about others’ thoughts and feelings (*28*). A recent meta-analysis of experimental fMRI studies on theory-of-mind and empathy found that TPJ was more strongly linked to mentalizing and moral cognition, than to (emotional) empathy (*11*).

Our data further suggested a mechanism for how dynamic concordance during pain anticipation led to pain relief for the patient. During pain, patients reporting the strongest analgesia also showed the strongest treatment-induced activation in a number of regions including the vlPFC (**Fig. S3**), which is implicated in both social mirroring (*29*) and psychosocially facilitated pain relief (*30*). We did not find that analgesia was associated with expectancy of treatment efficacy (Patients’ expectations: mean±SD: 4.00±2.80, r=−0.12, P=0.51; clinicians’ expectations: 5.52±2.53, r=−0.17, P=0.34), nor with brain responses in other regions related to expectancy-induced pain modulation, such as pgACC, dlPFC, and PAG (*31*, *32*). Instead, stronger patient analgesia was associated with more positive evaluations of the social interaction (e.g. the patient’s feeling of comfort from seeing the clinician). This may reflect potential differences in the brain circuitry responsible for socially mediated, relative to expectancy-mediated pain relief. Indeed, our results suggest a putative mechanism for social-context induced pain relief by which patients’ treatment-related vlPFC activation during pain statistically mediates the effect of anticipatory rTPJ concordance on analgesia.

A central question is why brain concordance and behavioral mirroring arises in the clinical encounter and is beneficial to patients. One possibility is that behavioral mirroring and synchrony may cause brain/physiological concordance, which, by promoting a positive affective-motivational state, leads to greater analgesia. From an evolutionary perspective, social affiliation signals support, care, and safety (*33*). One mode of this signaling may be behavioral synchronicity and neurobiological concordance, which are thought to support optimization of neural computation by reducing free energy and prediction errors (*34*), and thus represent a rewarding state associated with positive affect (*35*). The affective-motivational state induced by brain concordance may thus signal care and safety for the patient, reduce the perceived aversiveness/threat, and consequently the intensity of the painful stimulus during the clinical context. This would be consistent with two influential theoretical frameworks for understanding pain as a symptom. First, the ‘Motivation-decision model of pain’ posits that the brain continually makes (unconscious) decisions about the importance of nociceptive signals giving rise to pain, depending on the context (*36*). Second, the ‘Signaling theory of symptoms’ posits that besides promoting self-protection, a main function of clinical symptoms such as pain is *to motivate social signaling of the need for care* (*37*). Once this need is met, these symptoms should be attenuated. Hence, a positive clinical context characterized by high rapport, therapeutic alliance, and biobehavioral concordance, may serve as a safety signal for the patient; and consequently, the pain-evoking stimulus is deemed less salient, leading to analgesia. Recent studies have suggested behavioral mirroring and synchrony, e.g. in vocal acoustics (*16*), language style (*38*), posture (*17*), and gestures (*39*) as key features of clinical interactions. Here, we found that mirroring of facial expression was significantly associated with therapeutic alliance and analgesia.

Our study has several limitations. First, although we implemented a relatively naturalistic intake and consultation, and strived to maximize ecological validity during testing, the MRI environment necessitate the omission of several often-important psychosocial aspects in real-life therapeutic interactions (e.g. touch, sensitive proximity) (*40*). Future studies may address these aspects via analyses of cortical concordance using electroencephalography or near-infrared spectroscopy hyperscanning, allowing dyads to be recorded while interacting verbally and nonverbally in the same room. Second, there may be important aspects of the clinical relationship that develop over time and cannot be captured after a single intake. Indeed, while individual differences in analgesia were associated with social interaction quality, we did not find a mean group difference in analgesia between Clinical-Interaction and No-Interaction contexts. Future studies using a longitudinal design may elucidate how brain concordance and therapeutic alliance develops over time.

In conclusion, our study used a novel, comprehensive two-person approach to identify a putative brain-behavioral mechanism of the patient-clinician interaction. The findings represent an important first step toward specifying the non-specific components of the clinical encounter, and to establish the neuroscience supporting the patient-clinician relationship.

## Methods

### Subjects

Licensed acupuncturists were recruited from the local community and had completed at minimum a 3-year Masters-level program, or were in their final year of training and interning in clinics (Age: 44.32±12.81 (mean±SD), Race: 18 Caucasian, 1 Hispanic, 1 African-American, 1 Asian, 1 multiracial). Patients with chronic pain diagnosed with fibromyalgia (FM) for at least one year, meeting updated Wolfe et al. (*41*) criteria were recruited for the ‘Patient’ group (Age: 39.95±10.93, race: 18 Caucasian, 2 Hispanic, 2 African-American, 1 multiracial, all female). Clinicians ($150 per MRI session, $50 for non-MRI sessions) and patients ($100 per MRI session, $50 for non-MRI sessions) received monetary compensation for participation. The interval between the clinical intake and the Clinical-Interaction MRI was 8.32±14.07 days. The order was counterbalanced (which was limited by difficult scheduling logistics, Patients: 8 Clinical-Interaction first, 15 No-Interaction first. Clinicians: 14 Clinical-Interaction first, 8 No-Interaction first), and each participant contributed to two dyads, paired with a different partner, in order to avoid carryover effects related to the relationship. Thus, each dyad was unique. We scanned a total of 40 dyads, whereby we obtained complete MRI data from 37 dyads (19 Clinical-Interaction and 18 No-Interaction, with 2 dyads incomplete due to scanner malfunction, and 1 incomplete due to patient withdrawal due to claustrophobia mid-scan). Furthermore, three patients (1 ineligible, 2 due to scheduling issues) and 2 clinicians (due to scheduling issues) were enrolled but did not proceed to MRI scanning. Thus, 20 patients and 20 clinicians participated for at least 1 MRI visit, and were included in dyad-based analyses. Of these, 3 patients (2 due to scheduling issues, 1 due to claustrophobia) and 3 clinicians (2 due to scheduling issues, 1 due to scanner discomfort) dropped out after completing 1 MRI visit. Thus, for paired analyses, 17 female FM patients and 17 clinicians (12 female) completed both MRI visits. The Massachusetts General Hospital institutional review board approved the study, and all participants provided informed consent.

Since no relevant prior data existed on dynamic concordance, we could not estimate power using these dyad-based metrics. However, in our pilot data from clinicians providing treatment for the evoked pain of a ‘patient’ confederate (*12*), we observed a within-subject average Blood oxygen level-dependent (BOLD) percent change for ‘treatment’ minus ‘control’ (no treatment) of 1.25±1.53 (mean±SD). An a priori power analysis (paired, two-tailed, =0.05) indicated a minimum of 15 subjects (paired test) would be required for 85% power to detect this effect size (RStudio, function pwr.t.test, package pwr).

### Overall study protocol

Each patient came in for 3 or 4 visits, depending on whether they started with No-Interaction (4 visits) or Clinical-Interaction (3 visits – the initial consent/behavioral and clinical intake sessions were completed during the same visit). Each clinician came in for 3 visits – depending on interaction order, with their initial behavioral session completed on the same visit as the No-Interaction MRI (since clinicians’ initial behavioral session was shorter in duration than for patients), or just prior to the clinical intake session with the patient (for those starting with Clinical-Interaction). See below for further detail on each session.

#### Initial consent/behavioral session

After informed consent, participants were seated with a pressure cuff wrapped around their left lower leg, level with the gastrocnemius muscle. Participants went through a cuff pain calibration procedure to determine an individual stimulus intensity (pressure) level corresponding to moderate pain (40/100 pain rating). This pressure level was then used for all experimental cuff stimuli for this individual. Patients then had two acupuncture needles inserted on the anterior/distal aspect of the lower thigh, proximal to the cuff, with electrodes attached to each needle. Patients were then familiarized with the anticipation cue and pain stimuli, and received 6 cuff stimuli, 3 of which were preceded by a visual cue indicating that upcoming evoked pain would be treated with sub-sensory threshold electro-acupuncture (see below). For such ‘treatment’ trials, cuff pressure was surreptitiously reduced by 5, 10, and 20% of the target pressure (randomized order) in order to enhance expectations of treatment benefit, similar to boosting approaches previously used in investigations of the placebo effect (*32*, *42*).

#### Clinical intake

To maximize ecological validity, clinicians were instructed to perform a clinical consultation and intake with the patient ‘as similarly as possible to your daily practice’. Clinicians were not given restrictions on the duration of the intakes (mean±SD: 37:40±12:30 minutes:seconds, range: 21:32 – 54:40).

#### MRI sessions

Once the patient had been positioned in the MRI scanner (Skyra, 3T, Siemens Medical, Germany), the clinician entered the scanner room and led the patient through the process of acupuncture needling. MR-compatible titanium needles (0.22 mm thick, 40mm length, DongBang Acupuncture Inc. Boryeong, Korea) were inserted proximal to the cuff (2-3cm depth, acupoints ST-34 and SP-10), with MR-compatible electrodes attached to each needle. These acupoints were chosen for their local/segmental effects on a pain source delivered at the calf. Due to hospital policy, the actual needle penetration was performed by a staff acupuncturist with hospital credentials, but under direct supervision of the subject clinician, and evident to the patient. The clinician then attached MRI-compatible electrodes to the needles, and electrodes were connected to an electronic needle stimulation device (2Hz, 0.1mA, AS Super 4 Digital, Schwa-Medico, Wetzlar, Germany), controlled by the computer running the experimental protocol. The acupuncturist was then positioned in the other MRI scanner (Prisma, 3T, Siemens Medical, Germany), a 1-minute walk within the same building. In order to allow for unimpeded facial coverage for video transfer, both participants were positioned with an adapted coil configuration, using the 64-channel head coil bottom, and a small (4 channel) flex coil wrapped over the subjects’ forehead to cover the frontal lobes of the brain. Prior to the scan, participants were instructed that they would be free to communicate their feelings to the other person non-verbally using facial expressions, as long as they kept their head as still as possible. Prior to functional Magnetic Resonance Imaging (fMRI) scanning for the Clinical-Interaction session, the clinician was given the option to ‘check in’ with the patient via the between-scanner audio/video connection, in order to reinforce the clinical relationship.

### Self-report assessments

#### Therapeutic alliance

To assess the therapeutic empathy attributed to clinicians, patients filled out the validated Consultation And Relational Empathy (CARE) (*43*) scale after the intake and after each MRI visit, while clinicians filled out a modified CARE questionnaire with items phrased from the clinician’s point of view (*44*, *45*). Relational empathy was used as a proxy for therapeutic alliance.

#### Hyperscan Relationship Scale (HRS)

To assess ecological validity during MRI hyperscanning, as well as different qualities of the clinical interaction, we created a custom questionnaire to be filled out by patients (9 items, 2 reversed) and clinicians (10 items, 2 reversed) after each MRI visit (Visual Analog Scale, 0-10, anchors ‘Completely disagree’, ‘Completely agree’).

The patient scale included the following items: 1. I had frequent eye contact with the acupuncturist; 2. I felt as if the acupuncturist was in the same room as me; 3. I felt like I could communicate with the acupuncturist; 4. I felt comforted by seeing the acupuncturist; 5. I felt discomforted by seeing the acupuncturist; 6. I felt as if the acupuncturist was really trying to treat my leg pain with electroacupuncture; 7. The acupuncturist was genuinely concerned for me when I was in pain; 8. I expressed my feelings to the acupuncturist; 9. The acupuncturist was emotionally distant.

The clinician scale included the following items: 1. I had frequent eye contact with the patient; 2. I felt as if the patient was in the same room as me; 3. I felt like I could communicate with the patient; 4. I felt comforted by seeing the patient; 5. I felt discomforted by seeing the patient; 6. I thought my treatment was helping the patients pain; 7. I felt genuine concern for the patient when she was in pain; 8. I expressed my feelings to the patient; 9. I felt emotionally distant from the patient; 10. I cared whether I was providing electroacupuncture or not.

#### In-scanner ratings

At the end of each trial, participants used a MRI-compatible button-box to deliver 2 consecutive ratings (8 s each) on a Visual Analog Scale (VAS). Patients rated pain intensity (“How painful was the cuff?” with anchors “No pain” and “Most pain imaginable”), and affect related to either receiving treatment (“How did you feel about getting treated with electroacupuncture?” with anchors “Extremely negative” and “Extremely positive”) or not receiving treatment (“How did you feel about not getting the electroacupuncture?” with anchors “Extremely negative” and “Extremely positive”). Clinicians rated vicarious pain (“How painful was it for the patient?”), and affect related to either providing treatment (“How did you feel about doing the electroacupuncture?”) or not providing treatment (“How did you feel about not doing the electroacupuncture?”) with anchors “Extremely negative” and “Extremely positive”.

#### Treatment expectancy

Prior to the scan at each MRI visit, participants indicated their expectancy of electro-acupuncture treatment efficacy using a 0-10 VAS (Patient rating: “How much cuff pain relief do you expect to experience while being treated with electroacupuncture?” with anchors “No pain relief” to “Complete pain relief”; Clinician rating: “How much cuff pain relief do you expect the patient will experience while being treated with electroacupuncture?” with identical anchors).

### Other Materials

#### Cameras

For both participants, visual stimuli were projected onto a screen behind the MRI scanner bore, and participants viewed projected video through a mirror. To enable visual communication between the scanners, MRI-compatible cameras (Model 12M, MRC Systems GmbH, Heidelberg, Germany) were attached to the table-mounted mirror with each MRI scanner, and manually adjusted to capture the full face. The two-way video stream (20 Hz) was sent over a local network (measured to have consistent < 40 ms delay) and recorded for human facial expression artificial intelligence (AI) analyses (see below).

#### Microphones

MRI-compatible optical microphones Fibersound FOM1-MR, Micro Optics Technologies Inc., Cross Plains, WI, USA) were also set up in each MRI scanner to enable verbal communication *between* scans. To avoid speech-related motion *during* fMRI we decided to disallow verbal communication during fMRI scanning.

#### Software for stimulus presentation and signal synchronization

A custom in-house software (C++) was created for synchronizing fMRI scans between MRI scanners, transferring video and audio signals, and tracking the network delay between scanners. One laptop in each MRI scanner controlled the initiation of the fMRI scan acquisition sequence via remote trigger, the video stream, the experimental design visual stimuli, onset/offset of the cuff stimuli via remote trigger, and recording of in-scanner ratings. Both laptops were connected through a Local Area Network. The MRI teams in each control room communicated with one another via phone and, when ready to start, the master computer (patient MRI control room) sent a signal to the slave computer (clinician MRI control room) to initiate the fMRI pulse sequence. Thus, after a lag corresponding to the current network delay (mean±SD=81.6±38.1 milliseconds, calculated as mean of 10 network pings), each computer initiated the fMRI pulse sequence locally. This procedure ensured synchronized timing of the two fMRI time series, video streams, and experimental protocols.

### Statistical analysis

All non-imaging statistical analyses were completed using R (RStudio 1.1.456) and JASP (version 0.10, Jasp Team, Amsterdam, Netherlands). Threshold for statistical significance was set at alpha=0.05.

#### Therapeutic alliance

To evaluate whether therapeutic alliance (CARE score) was different between sessions, we performed separate one-way repeated measures ANOVAs for the patient group and the clinician group, each with three levels (Intake, Clinical-Interaction MRI, and No-Interaction MRI). We then performed follow-up contrasts comparing the different sessions.

#### Influence of therapeutic alliance at the intake on social interaction at the MRI session

In order to evaluate whether the relationship established during the intake carried over to the Clinical-Interaction MRI, we performed two ANCOVAs (separately for patient-rated and clinician-rated scores), with HRS values at MRI (see Hyperscan relationship scale above) as the dependent variable, as an indicator of social interaction quality. Therapeutic alliance at intake (CARE_Intake_) was used as a continuous predictor and HRS Item was used as a categorical predictor to investigate potential differences between items of the HRS scale.

#### Pain and affect

Ratings of cuff pain intensity and affect were analyzed using separate repeated measures ANOVAs with factors ‘Treatment condition’ (Treatment, No-Treatment), ‘Clinical context’ ‘(Clinical-Interaction, No-Interaction), and ‘Order’ as a between-subjects factor (Clinical-Interaction first, No-Interaction first).

#### Association between social relationship and patient analgesia

To evaluate whether differences in the social interaction between dyads were associated with analgesia, we performed an ANCOVA with Analgesia (Pain_Treat-NoTreat_) as the dependent variable, HRS values (Patient-rated) as a continuous predictor and ‘Clinical context’ (Clinical-Interaction, No-Interaction) and HRS Item (See ‘Hyperscan Relationship Scale’ above) as categorical predictors.

#### Treatment expectancy

To evaluate whether prior expectancy of therapeutic efficacy predicted treatment-related pain relief, we calculated Pearson’s correlation coefficients between expectancy as rated by the patient and the clinician prior to scanning Vs. analgesia (mean Pain_Treat-NoTreat_) during scanning.

#### Facial expression analyses

Facial expressions during fMRI scanning were analyzed using automated facial feature extraction (Affectiva, Cambridge, MA). The Affectiva Facial Expression Analysis algorithm is based on the Emotional Facial Action Coding System (*46*) and trained on ~8 million images and videos of faces. Due to limited field-of-view in forehead and chin regions for some participants, we were able to fully analyze patient data from 24 dyads and clinician data from 21 dyads (17 dyads had adequate data for both patient and clinician data). For the Affectiva algorithm, 33 facial landmarks are initially identified, which were used to estimate 21 facial action units. These units were then mapped onto 7 basic emotional expressions (joy, fear, disgust, sadness, anger, surprise, contempt) and 2 core expressions (valence, engagement). We calculated these 9 expressions frame-by-frame and averaged across each trial duration (separately for anticipation and pain/treatment phases).

Overall mirroring: Behavioral mimicry such as the mirroring of facial expressions is thought to be fundamental for social development (*47*, *48*), and a cornerstone of the establishment and maintenance of human bonds (*49*, *50*), including in the patient-clinician interaction (*51*). As the specific facial expressions mirrored can be variable across individuals and interactions, we decided to investigate correspondence within each dyad and across different expressions. We first calculated the difference in each expression between anticipation of Pain/Treatment relative to Pain/No-treatment. Using these difference scores, we then calculated a Pearson’s correlation coefficient between the patient and the clinician of each dyad. This coefficient was then Fisher’s R-to-Z transformed and used as a metric of each dyad’s overall facial mirroring. We then investigated if facial mirroring was associated with therapeutic alliance (CARE scores) and analgesia.

#### fMRI analysis

##### Treatment-related differences in pain-related brain activation

Details on MRI acquisition and preprocessing are described in the Supplementary Methods. For all whole-brain group fMRI analyses, significance testing was performed using FSL FLAME 1+2 with cluster correction for multiple comparisons (z=2.3, α=0.05) (*52*). In order to investigate treatment-related differences in brain response during pain, we first performed single-subject first-level GLM analyses using FILM with local autocorrelation correction (*53*). For each of the 2 runs (6 trials each) we modeled periods corresponding to pain stimulation (Treat, NoTreat) as regressors. In the same design matrix, we also modeled ratings periods and the 6 motion parameter time series as regressors of no interest. We computed 2 bi-directional contrasts: Pain_Treat_-Rest, Pain_NoTreat_-Rest. In second-level fixed-effects analyses, we averaged these contrast parameter estimates across both runs and both visits (Clinical-Interaction and No-Interaction) for each patient. The resulting contrast parameter estimate maps were then passed up to a group analysis where a whole-brain group mean was calculated for all patients.

##### Regression with analgesia

To investigate brain regions where treatment-related change in BOLD contrast correlated with analgesia, we performed a whole-brain regression GLM using each patient’s mean analgesia (Pain_Treat_ – Pain_NoTreat_) ratings as a regressor of interest.

##### Overall brain response to anticipation and pain

Patient-clinician concordance related to therapeutic alliance and pain outcomes could be driven by social interaction during the anticipation of both treated and non-treated pain. Therefore, we first calculated overall brain response to anticipation of pain irrespective of Treat/NoTreat conditions, followed by a group conjunction between patients and clinicians, to identify Regions of Interest (ROI) for concordance analyses.

Single-subject GLM analyses were performed using FILM with local autocorrelation correction. Similar to above, for each of the 2 runs we modeled periods corresponding to anticipation of pain and pain stimulation as regressors. We also modeled ratings periods and the 6 motion parameter time series as regressors of no interest. We computed bi-directional contrasts for Anticipation-Rest and Pain-Rest. In second-level fixed-effects analyses, we averaged these contrast parameter estimates across both runs and both visits (Clinical-Interaction and No-Interaction) for each individual. We then passed the resulting contrast parameter estimate maps up to group analyses (separately for patients and clinicians), indicating overall response to 1) anticipation of pain (patients) and preparing to treat/not treat (clinicians); and 2) pain (patients) and observing pain and treating/not treating (clinicians). In order to identify shared activation between patients and clinicians during the anticipation phase, for ROI identification for concordance analyses, we first performed a conjunction of the minimum statistic between these two maps. This group conjunction map was corrected for multiple comparisons using False Discovery Rate (α=0.05). We then intersected this whole-brain map with *a priori* structural ROIs involved in social mirroring, empathy, and theory-of-mind (see *a priori* ROIs section), to yield more specific functional ROIs for use in concordance analyses.

##### Patient-clinician dynamic concordance in brain activity

To assess dynamic concordance in brain activity between patients and clinicians, we first performed two first-level GLMs (one for each fMRI scan run), with each trial (anticipation period) as a separate regressor (**Fig. S5**). We also modeled each pain period as a separate regressor of no interest. This produced a total of 12 pain anticipation parameter estimate maps (across both runs) for each individual. We then extracted the mean Zstat value from each individual’s right Temporoparietal Junction (rTPJ) for each of the 12 anticipation trials, as defined by the group conjunction map intersected with the anatomical ROI. For each dyad, we performed a second-level whole-brain regression analysis of the clinician’s brain, using the trial-by-trial rTPJ Zstats from the patient as a regressor, and vice versa. Thus, we obtained a whole-brain map for each individual showing regions *dynamically* concordant (across trials) with the dynamics of the partner’s rTPJ response throughout the interaction. Next, we performed a whole-brain group contrast between Clinical-Interaction and No-Interaction to investigate regions where dynamic concordance was increased by Clinical-Interaction.

##### Mediation analysis

Finally, we explored whether treatment-induced change in patients’ brain response during pain reflecting analgesia (i.e. Treat-NoTreat) mediated the influence of brain concordance on analgesia ratings. We decided to focus on rTPJ-to-rTPJ concordance, as this metric was correlated with analgesia. We first extracted the mean Zstat from the rTPJ region of each clinician’s whole-brain concordance map with the patient’s rTPJ, as a metric of each dyad’s rTPJ-rTPJ concordance, which was then used as the independent variable. For the mediator variable, we focused on the ventrolateral Prefrontal Cortex (vlPFC), as this region is both a key region for social mirroring (*12*, *54*– *57*), and has been implicated in psychosocial and placebo analgesia (*30*, *58*–*61*). The vlPFC ROI was chosen based on an intersection between the Pain_Treat-NoTreat_ regression map and an anatomical mask (Inferior Frontal Gyrus, pars triangularis and pars opercularis, combined mask, p(>30%). We extracted the mean Zstat value from the vlPFC_Treat-NoTreat_ Zstat ROI from each patient, which we then used as a mediator variable in further analyses. Additionally, since the Pain_Treat-NoTreat_ contrast was also increased for patients in brain regions beyond vlPFC (e.g. TPJ, dorsolateral PFC (dlPFC), Superior Temporal Sulcus (STS), and medial PFC (mPFC)), we also explored whether these regions mediated the association between concordance and analgesia. Each patient’s (Pain_Treat-NoTreat_) rating difference was used as the dependent variable. We used the R package ‘Mediation’ for mediation analyses (*62*). We tested for statistical significance using a boot strapping approach (1000 iterations, α=0.05), and considered the mediation significant if the total indirect effect (a*b) was statistically significant, while the previously significant direct effect (path c) became non-significant after controlling for the mediator (c’) (*63*).

## Acknowledgments

We thank Andre van der Kouwe, Thomas Witzel, Paul Wighton, Matthew Rosen, and Simon Sigalovsky for technical assistance setting up the fMRI hyperscanning environment, Marco Loggia for helpful input on research design, and Katie Walker for help with acupuncture administration.

## References

1. R. Grob, G. Darien, D. Meyers, Why Physicians Should Trust in Patients. JAMA. 321, 1347 (2019).

2. K. Whetten, J. Leserman, R. Whetten, J. Ostermann, N. Thielman, M. Swartz, D. Stangl, Exploring Lack of Trust in Care Providers and the Government as a Barrier to Health Service Use. Am J Public Health. 96, 716–721 (2006).

3. T.J. Kaptchuk, J.M. Kelley, L.A. Conboy, R.B. Davis, C.E. Kerr, E.E. Jacobson, I. Kirsch, R.N. Schyner, B.H. Nam, L.T. Nguyen, M. Park, A. L. Rivers, C. McManus, E. Kokkotou, D.A. Drossman, P. Goldman, A.J. Lembo, Components of placebo effect: randomised controlled trial in patients with irritable bowel syndrome. BMJ. 336, 999–1003 (2008).

4. P.H. Ferreira, M.L. Ferreira, C.G. Maher, K.M. Refshauge, J. Latimer, R. D. Adams, The Therapeutic Alliance Between Clinicians and Patients Predicts Outcome in Chronic Low Back Pain. Physical Therapy. 93, 470–478 (2013).

5. M. E. Suarez-Almazor, C. Looney, Y. Liu, V. Cox, K. Pietz, D.M. Marcus, R.L. Street, A randomized controlled trial of acupuncture for osteoarthritis of the knee: Effects of patient-provider communication. Arthritis Care & Research. 62, 1229–1236 (2010).

6. L. Dyche, D. Swiderski, The effect of physician solicitation approaches on ability to identify patient concerns. J GEN INTERN MED. 20, 267–270 (2005).

7. A.H. Kamal, J.H. Bull, S.P. Wolf, K.M. Swetz, T.D. Shanafelt, K. Ast, D. Kavalieratos, C.T. Sinclair, A.P. Abernethy, Prevalence and Predictors of Burnout Among Hospice and Palliative Care Clinicians in the U.S. Journal of Pain and Symptom Management. 51, 690–696 (2016).

8. D. Musa, R. Schulz, R. Harris, M. Silverman, S.B. Thomas, Trust in the Health Care System and the Use of Preventive Health Services by Older Black and White Adults. Am J Public Health. 99, 1293–1299 (2009).

9. H. Leis, P. Garg, I. Soh, “Right Place, Right Time: Marketplace Responses to the Health Information Needs of Vulnerable Consumers: Final Report” (2017), (available at https://owy.mn/3cdsuRo).

10. T.J. Kaptchuk, F.G. Miller, Placebo Effects in Medicine. New England Journal of Medicine. 373, 8–9 (2015).

11. D. Bzdok, L. Schilbach, K. Vogeley, K. Schneider, A.R. Laird, R. Langner, S.B. Eickhoff, Parsing the neural correlates of moral cognition: ALE meta-analysis on morality, theory of mind, and empathy. Brain Struct Funct. 217, 783–796 (2012).

12. K.B. Jensen, P. Petrovic, C.E. Kerr, I. Kirsch, J. Raicek, A. Cheetham, R. Spaeth, A. Cook, R.L. Gollub, J. Kong, T.J. Kaptchuk, Sharing pain and relief: neural correlates of physicians during treatment of patients. Molecular Psychiatry. 19, 392–398 (2014).

13. E. Redcay, L. Schilbach, Using second-person neuroscience to elucidate the mechanisms of social interaction. Nat Rev Neurosci. 20, 495–505 (2019).

14. J. Levy, R. Feldman, Synchronous Interactions Foster Empathy. J Exp Neurosci. 13, 1179069519865799 (2019).

15. U. Hasson, C.D. Frith, Mirroring and beyond: coupled dynamics as a generalized framework for modelling social interactions. Phil. Trans. R. Soc. B. 371, 20150366 (2016).

16. Z.E. Imel, J.S. Barco, H.J. Brown, B.R. Baucom, J.S. Baer, J.C. Kircher, D.C. Atkins, The association of therapist empathy and synchrony in vocally encoded arousal. Journal of Counseling Psychology. 61, 146–153 (2014).

17. F. Ramseyer, W. Tschacher, Nonverbal synchrony in psychotherapy: Coordinated body movement reflects relationship quality and outcome. Journal of Consulting and Clinical Psychology. 79, 284–295 (2011).

18. A. Finset, K. Ørnes, Empathy in the Clinician–Patient Relationship: The Role of Reciprocal Adjustments and Processes of Synchrony. Journal of Patient Experience. 4, 64–68 (2017).

19. C.D. Marci, J. Ham, E. Moran, S.P. Orr, Physiologic Correlates of Perceived Therapist Empathy and Social-Emotional Process During Psychotherapy: The Journal of Nervous and Mental Disease. 195, 103–111 (2007).

20. M.B. Schippers, A. Roebroeck, R. Renken, L. Nanetti, C. Keysers, Mapping the information flow from one brain to another during gestural communication. P Natl Acad Sci USA. 107, 9388–9393 (2010).

21. L.J. Silbert, C.J. Honey, E. Simony, D. Poeppel, U. Hasson, Coupled neural systems underlie the production and comprehension of naturalistic narrative speech. Proc Natl Acad Sci U S A. 111, E4687–96 (2014).

22. G.J. Stephens, L.J. Silbert, U. Hasson, Speaker-listener neural coupling underlies successful communication. Proceedings of the National Academy of Sciences. 107, 14425–14430 (2010).

23. E. Bilek, M. Ruf, A. Schäfer, C. Akdeniz, V.D. Calhoun, C. Schmahl, C. Demanuele, H. Tost, P. Kirsch, A. Meyer-Lindenberg, Information flow between interacting human brains: Identification, validation, and relationship to social expertise. Proceedings of the National Academy of Sciences. 112, 5207–5212 (2015).

24. S.W. Mercer, M. Maxwell, D. Heaney, G.C. Watt, The consultation and relational empathy (CARE) measure: development and preliminary validation and reliability of an empathy-based consultation process measure. Family practice. 21, 699–705 (2004).

25. A.P. Burgess, On the interpretation of synchronization in EEG hyperscanning studies: a cautionary note. Front. Hum. Neurosci. 7 (2013), doi:10.3389/fnhum.2013.00881.

26. S.L. Koole, W. Tschacher, Synchrony in Psychotherapy: A Review and an Integrative Framework for the Therapeutic Alliance. Front. Psychol. 7 (2016), doi:10.3389/fpsyg.2016.00862.

27. D.A. Matthews, Making Connexions: Enhancing the Therapeutic Potential of PatientClinician Relationships. Ann Intern Med. 118, 973 (1993).

28. M. Schurz, M.G. Tholen, J. Perner, R.B. Mars, J. Sallet, Specifying the brain anatomy underlying temporo-parietal junction activations for theory of mind: A review using probabilistic atlases from different imaging modalities. Hum Brain Mapp. 38, 4788–4805 (2017).

29. S. G. Shamay-Tsoory, J. Aharon-Peretz, D. Perry, Two systems for empathy: a double dissociation between emotional and cognitive empathy in inferior frontal gyrus versus ventromedial prefrontal lesions. Brain. 132, 617–27 (2009).

30. L.Y. Atlas, T.D. Wager, in Placebo, F. Benedetti, P. Enck, E. Frisaldi, M. Schedlowski, Eds. (Springer Berlin Heidelberg, Berlin, Heidelberg, 2014; http://link.springer.com/10.1007/978-3-662-44519-8_3), vol. 225, pp. 37–69.

31. A. Tinnermann, S. Geuter, C. Sprenger, J. Finsterbusch, C. Büchel, Interactions between brain and spinal cord mediate value effects in nocebo hyperalgesia. Science. 358, 105–108 (2017).

32. F. Eippert, U. Bingel, E.D. Schoell, J. Yacubian, R. Klinger, J. Lorenz, C. Büchel, Activation of the Opioidergic Descending Pain Control System Underlies Placebo Analgesia. Neuron. 63, 533–543 (2009).

33. J. Panksepp, J.B. Panksepp, Toward a cross-species understanding of empathy. Trends in Neurosciences. 36, 489–496 (2013).

34. L. Koban, A. Ramamoorthy, I. Konvalinka, Why do we fall into sync with others? Interpersonal synchronization and the brain’s optimization principle. Social Neuroscience. 14, 1–9 (2019).

35. L. Beckes, J.A. Coan, Social Baseline Theory: The Role of Social Proximity in Emotion and Economy of Action. Social and Personality Psychology Compass. 5, 976–988 (2011).

36. H.L. Fields, Understanding how opioids contribute to reward and analgesia. Region Anesth Pain M. 32, 242–246 (2007).

37. L. Steinkopf, The Signaling Theory of Symptoms: An Evolutionary Explanation of the Placebo Effect. Evolutionary Psychology. 13, 147470491560055 (2015).

38. S.P. Lord, E. Sheng, Z.E. Imel, J. Baer, D.C. Atkins, More Than Reflections: Empathy in Motivational Interviewing Includes Language Style Synchrony Between Therapist and Client. Behavior Therapy. 46, 296–303 (2015).

39. W. Tschacher, G.M. Rees, F. Ramseyer, Nonverbal synchrony and affect in dyadic interactions. Front. Psychol. 5, 1323 (2014).

40. M.L. Knapp, J.A. Hall, Nonverbal communication in human interaction (4th ed.) (Fort Worth, TX: Harcourt Brace College, 1997).

41. F. Wolfe, W. Häuser, Fibromyalgia diagnosis and diagnostic criteria. Annals of Medicine. 43, 495–502 (2011).

42. S. T. Au Yeung, B. Colagiuri, P.F. Lovibond, L. Colloca, Partial reinforcement, extinction, and placebo analgesia. Pain. 155, 1110–7 (2014).

43. S.W. Mercer, M. Maxwell, D. Heaney, G.C. Watt, The consultation and relational empathy (CARE) measure: development and preliminary validation and reliability of an empathy-based consultation process measure. Fam Pract. 21, 699–705 (2004).

44. M. Phillips, A. Lorie, J. Kelley, S. Gray, H. Riess, Long-term effects of empathy training in surgery residents: a one year follow-up study. European Journal for Person Centered Healthcare. 1, 326–332 (2013).

45. H. Riess, J.M. Kelley, R. Bailey, P.M. Konowitz, S.T. Gray, Improving Empathy and Relational Skills in Otolaryngology Residents: A Pilot Study. Otolaryngol Head Neck Surg. 144, 120–122 (2011).

46. W. Friesen, P. Ekman, EMFACS-7: Emotional facial action coding system (1983).

47. A.N. Meltzoff, M.K. Moore, Imitation of facial and manual gestures by human neonates. Science. 198, 74–78 (1977).

48. H. Rayson, J.J. Bonaiuto, P.F. Ferrari, L. Murray, Early maternal mirroring predicts infant motor system activation during facial expression observation. Scientific Reports. 7, 11738 (2017).

49. T.L. Chartrand, J.A. Bargh, The chameleon effect: The perception– behavior link and social interaction. Journal of Personality and Social Psychology. 76, 893 (19990701).

50. M. Salazar Kämpf, H. Liebermann, R. Kerschreiter, S. Krause, S. Nestler, S. C. Schmukle, Disentangling the Sources of Mimicry: Social Relations Analyses of the Link Between Mimicry and Liking. Psychol Sci. 29, 131–138 (2018).

51. A. Finset, T.A. Mjaaland, The medicalconsultation viewed asa value chain: A neurobehavioral approach to emotion regulation in doctor–patient interaction. Patient Education and Counseling. 74, 323–330 (2009).

52. M.W. Woolrich, T. E.J. Behrens, C.F. Beckmann, M. Jenkinson, S.M. Smith, Multilevel linear modelling for FMRI group analysis using Bayesian inference. NeuroImage. 21, 1732–1747 (2004).

53. M.W. Woolrich, B.D. Ripley, M. Brady, S.M. Smith, Temporal autocorrelation in univariate linear modeling of FMRI data. NeuroImage. 14, 1370–86 (2001).

54. E. A.R. Losin, C.-W. Woo, A. Krishnan, T.D. Wager, M. Iacoboni, M. Dapretto, Brain and psychological mediators of imitation: sociocultural versus physical traits. Culture and Brain. 3, 93–111 (2015).

55. C. Becchio, A. Cavallo, C. Begliomini, L. Sartori, G. Feltrin, U. Castiello, Social grasping: from mirroring to mentalizing. Neuroimage. 61, 240–8 (2012).

56. L. Budell, P. Jackson, P. Rainville, Brain responses to facial expressions of pain: emotional or motor mirroring? Neuroimage. 53, 355–63 (2010).

57. L. Budell, M. Kunz, P.L. Jackson, P. Rainville, Mirroring pain in the brain: emotional expression versus motor imitation. PLoS One. 10, e0107526 (2015).

58. E. Vachon-Presseau, Brain and psychological determinants of placebo pill response in chronic pain patients. NATURE COMMUNICATIONS. 9, 1–15 (2018).

59. I. Tracey, Getting the pain you expect: mechanisms of placebo, nocebo and reappraisal effects in humans. nature medicine. 16, 1277–1283 (2010).

60. M. Amanzio, F. Benedetti, C.A. Porro, S. Palermo, F. Cauda, Activation likelihood estimation meta-analysis of brain correlates of placebo analgesia in human experimental pain. Hum. Brain Mapp. 34, 738–752 (2013).

61. M.D. Lieberman, J.M. Jarcho, S. Berman, B.D. Naliboff, B.Y. Suyenobu, M. Mandelkern, E.A. Mayer, The neural correlates of placebo effects: a disruption account. NeuroImage. 22, 447–455 (2004).

62. D. Tingley, T. Yamamoto, K. Hirose, L. Keele, K. Imai, mediation: R package for causal mediation analysis. UCLA Statistics/American Statistical Association (2014) (available at https://dspace.mit.edu/handle/1721.1/91154).

63. K. Imai, L. Keele, D. Tingley, A general approach to causal mediation analysis. Psychological Methods. 15, 309–334 (2010).

