## Supplementary Information for "Dynamic brain-to-brain concordance and behavioral mirroring as a mechanism of the patient-clinician interaction"

### Supplementary Methods and Materials

#### Stimuli

##### *Evoked cuff pain*

Deep-tissue pain was applied using the Hokanson Rapid Cuff Inflator (D. E. Hokanson, Inc., Bellevue, WA, USA). Compared to cutaneous quantitative sensory testing (QST) techniques (e.g., contact heat), deep sustained pain better mimics clinical pain (*1*, *2*), thus providing a more clinically-relevant measure. Unlike more superficial methods of evaluating mechanical sensitivity, cuff pain responses are only marginally affected by sensitization or desensitization of the skin, indicating that this procedure primarily assesses sensitivity in muscle and other deep tissues (*3*–*6*). Such cuff pressure algometry is a recently characterized method that is now included in many QST evaluations (*3*, *4*, *7*). We have considerable experience applying these techniques in chronic pain patients, with and without neuroimaging, and have found that even subjects with severe fibromyalgia are generally able to tolerate these procedures without any lasting discomfort (*8*, *9*). The cuff was attached to the patient's left lower leg prior to scanning and was inflated for 15 s duration trials during the experiment.

##### *Electroacupuncture*

Our decision to use electroacupuncture applied by acupuncturists for evoked cuff pain as an experimental model for dyadic interaction with pain therapy was motivated by our aim to investigate both a consultation/intake and actual treatment with measurable pain outcomes. As opposed to many other pain therapies, this electroacupuncture/cuff pain model enabled us to incorporate acute pain therapy in the MRI environment, with acupuncturists remotely applying treatment relevant to their clinical practice, thus increasing ecological validity while maintaining experimental control.

Prior to experimental testing, patients had acupuncture needles (0.20 mm diameter, 25 cm length, Asiamed gauge) inserted with approximately 2-3cm depth at acupoints ST-34 and SP-10 on the anterior/distal aspect of the lower thigh, proximal to the cuff, with electrodes attached to each needle. While we were interested in the influence of therapeutic alliance, and not specific acupuncture effects, on pain outcomes, we wanted to avoid the need for authorized deception in clinicians' IRB consent form. We therefore included trials with both verum and sham electro-acupuncture in a double-blind manner. We used a minimal sub-sensory threshold level (0.1 mA) for verum trials in order to avoid unblinding patients due to any sensory feedback from the electrical stimulation. Thus, verum/sham electro-acupuncture was administered in a double-blind manner, in a pseudorandomized order across trials. Clinicians were instructed to press-and-hold one button for applying treatment, and another ("inactive") button for No-treatment.

We did not hypothesize differences in pain for verum vs sham electro-acupuncture treatment, and indeed, a paired t-test comparing patient-rated pain intensity between verum and sham trials did not suggest a difference in pain intensity (t=0.83, P=0.42). Consequently, we pooled verum/sham trials as 'Treatment' in further statistical analyses, and interpreted intra-individual differences between 'Treatment' and 'No-Treatment' trials as psychosocially induced pain relief ('analgesia').

#### MRI acquisition and preprocessing

*MRI acquisition*

Blood oxygen level-dependent (BOLD) fMRI data were collected from each scanner (Patient scanner: Siemens 3T Skyra; Clinician scanner: Siemens 3T Prisma) using a whole brain, simultaneous multi-slice, T2*-weighted gradient echo BOLD echo-planar imaging pulse sequence (repetition time = 1250 ms, echo time = 33 ms, flip angle = 65°, voxel size = 2 cm isotropic, number of slices = 75, Simultaneous Multi-Slice factor = 5). A high-resolution structural volume (multi-echo MPRAGE) was collected to facilitate anatomical localization and spatial registration of individual fMRI-BOLD volumes to MNI152 standard space (repetition time = 2530 ms, echo time = 1.69 ms, flip angle = 7°, voxel size = 1 mm isotropic). Importantly, to enable a full-face view of each participant for better facial expression tracking by research subjects and the facial expression digitization software (see below), we combined the occipital/bottom portion of a 64 Channel head coil with a flex coil (4 channel) attached to the forehead.

*fMRI preprocessing*

Preprocessing of individual fMRI datasets was carried out using tools from FMRIB's Software Library (FSL, v6.0.0; www.fmrib.ox.ac.uk/fsl), and included the following steps: slice-timing correction, motion correction (MCFLIRT) (*10*), correction of spatial inhomogeneity (TOPUP) (*11*, *12*), nonbrain tissue removal (BET) (*13*), spatial smoothing (full width at half maximum = 4mm), temporal high-pass filtering (f=0.011 Hz as computed by FSL's cutoffcalc), and grand-mean intensity normalization by a single multiplicative factor. For each subject, both runs were realigned (6 degrees of freedom) to a common space (7th volume of the first run) before the first-level general linear model (GLM) analyses. The transformation matrix for registration between functional and high-resolution anatomical volumes was calculated using Boundary Based Registration (bbregister, Freesurfer, v6.0.0 (*14*)). Two participants had one of their two fMRI runs excluded from analysis due to excessive head motion, based on the following exclusion criteria: 1) >2° head rotation in any direction, and 2) >2 mm frame-by-frame displacement. After excluding these data, mean head rotation was 0.05±0.02 (mean±SD) and mean frame-by-frame displacement was 0.13±0.05. An unpaired t-test indicated higher frame-by-frame displacement for patients (0.15±0.05) relative to clinicians (0.11±0.04, t=4.32, P<0.001), but there was no significant group difference for rotation (t=1.74, P=0.09). For registration from structural to standard space (Montreal Neurological Institute, MNI, 152), we used FSL's Linear registration tool (FLIRT, 12 degrees of freedom) (*10*, *15*), followed by FSL's non-linear registration tool (FNIRT) (*16*). All single-subject analyses were performed in functional space, and then registered to MNI152 standard space before dyadic and group analyses.

### References


1. P. Rainville, J. S. Feine, M. C. Bushnell, G. H. Duncan, A Psychophysical Comparison of Sensory and Affective Responses to Four Modalities of Experimental Pain. *Somatosensory & Motor Research*. **9**, 265–277 (1992).
2. M. Curatolo, L. Arendt-Nielsen, S. Petersen-Felix, Central Hypersensitivity in Chronic Pain: Mechanisms and Clinical Implications. *Physical Medicine and Rehabilitation Clinics*. **17**, 287–302 (2006).
3. R. Polianskis, T. Graven-Nielsen, L. Arendt-Nielsen, Spatial and temporal aspects of deep tissue pain assessed by cuff algometry. *Pain*. **100**, 19–26 (2002).
4. R. Polianskis, T. Graven-Nielsen, L. Arendt-Nielsen, Pressure-pain function in desensitized and hypersensitized muscle and skin assessed by cuff algometry. *The Journal of Pain*. **3**, 28–37 (2002).
5. R. Polianskis, T. Graven-Nielsen, L. Arendt-Nielsen, Modality-specific facilitation and adaptation to painful tonic stimulation in humans. *European Journal of Pain*. **6**, 475–484 (2002).

6. R. Polianskis, T. Graven-Nielsen, L. Arendt-Nielsen, Computer-controlled pneumatic pressure algometry—a new technique for quantitative sensory testing. *European Journal of Pain*. **5**, 267–277 (2001).

7. V. Napadow, R. R. Edwards, C. M. Cahalan, G. Mensing, S. Greenbaum, A. Valovska, A. Li, J. Kim, Y. Maeda, K. Park, A. D. Wasan, Evoked Pain Analgesia in Chronic Pelvic Pain Patients Using Respiratory-Gated Auricular Vagal Afferent Nerve Stimulation. *Pain Med*. **13**, 777–789 (2012).

8. M. L. Loggia, C. Berna, J. Kim, C. M. Cahalan, R. L. Gollub, A. D. Wasan, R. E. Harris, R. R. Edwards, V. Napadow, Disrupted brain circuitry for pain-related reward/punishment in fibromyalgia. *Arthritis Rheumatol*. **66**, 203–12 (2014).

9. J. Kim, M. L. Loggia, C. M. Cahalan, R. E. Harris, F. Beissner, R. G. Garcia, H. Kim, R. Barbieri, A. D. Wasan, R. R. Edwards, V. Napadow, The somatosensory link in fibromyalgia: functional connectivity of the primary somatosensory cortex is altered by sustained pain and is associated with clinical/autonomic dysfunction. *Arthritis Rheumatol*. **67**, 1395–405 (2015).

10. M. Jenkinson, P. Bannister, M. Brady, S. Smith, Improved optimization for the robust and accurate linear registration and motion correction of brain images. *Neuroimage*. **17**, 825–841 (2002).

11. J. L. R. Andersson, S. Skare, J. Ashburner, How to correct susceptibility distortions in spin-echo echo-planar images: application to diffusion tensor imaging. *Neuroimage*. **20**, 870–888 (2003).

12. S. M. Smith, M. Jenkinson, M. W. Woolrich, C. F. Beckmann, T. E. J. Behrens, H. Johansen-Berg, P. R. Bannister, M. De Luca, I. Drobnjak, D. E. Flitney, R. K. Niazy, J. Saunders, J. Vickers, Y. Y. Zhang, N. De Stefano, J. M. Brady, P. M. Matthews, Advances in functional and structural MR image analysis and implementation as FSL. *Neuroimage*. **23**, S208–S219 (2004).

13. S. M. Smith, Fast robust automated brain extraction. *Human brain mapping*. **17**, 143–55 (2002).

14. D. N. Greve, B. Fischl, Accurate and robust brain image alignment using boundary-based registration. *Neuroimage*. **48**, 63–72 (2009).

15. M. Jenkinson, S. Smith, A global optimisation method for robust affine registration of brain images. *Medical image analysis*. **5**, 143–56 (2001).

16. J. Andersson, M. Jenkinson, S. Smith, "Non-linear registration, aka spatial normalisation, FMRIB technical report TR07JA2" (2010).

17. M. Zunhammer, U. Bingel, T. D. Wager, C. Placebo Imaging, Placebo Effects on the Neurologic Pain Signature: A Meta-analysis of Individual Participant Functional Magnetic Resonance Imaging Data. *JAMA Neurol*. **75**, 1321–1330 (2018).

18. T. D. Wager, L. Y. Atlas, M. A. Lindquist, M. Roy, C.-W. Woo, E. Kross, An fMRI-Based Neurologic Signature of Physical Pain. *New England Journal of Medicine*. **368**, 1388–1397 (2013).

### Supplementary Figures

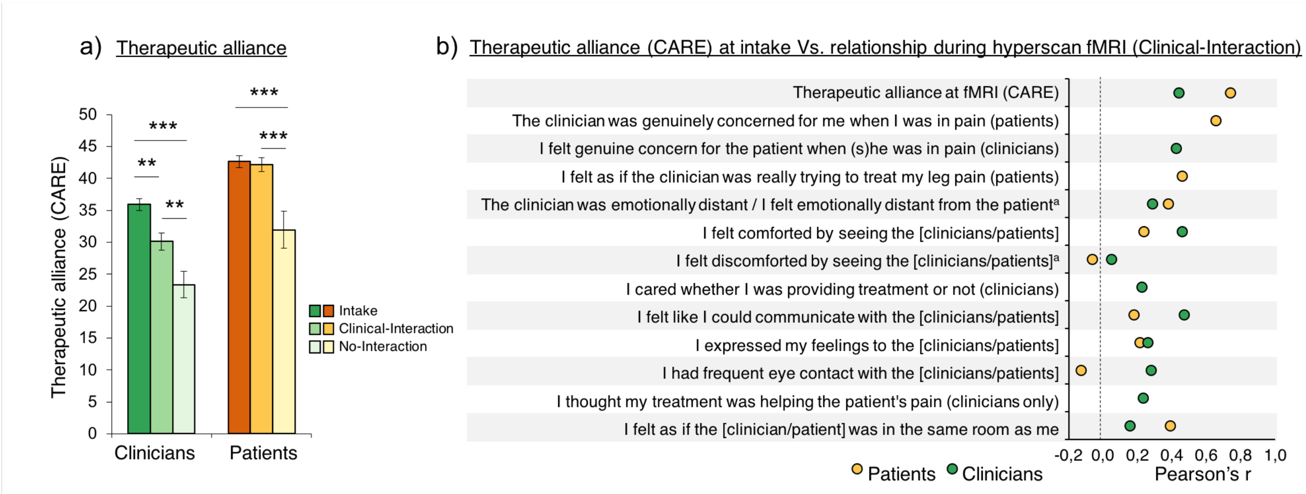


**Fig. S1 | Self-reported patient-clinician therapeutic alliance (CARE) across sessions.** a) Similarly to patients (see main Article), clinicians reported different levels of therapeutic alliance (CARE scores) depending of the context of the dyadic clinical interaction (F(1.82,25.42)=12.83, P<0.001, ηp2=0.48). Specifically, the No-interaction MRI context (mean±SD=23.56±7.94) was rated lower than both Intake (35.69±4.08, t=5.95, P<0.001, Cohen's d=1.49, 95% Confidence Interval [CI]=6.63,17.62) and Clinical-Interaction MRI (30.25±5.89, t=2.84, P=0.01, d=0.71, CI=0.34,13.03) contexts. Furthermore, therapeutic alliance at Intake was rated higher than at Clinical-Interaction MRI (t=3.11, P=0.01, d=0.78, CI=0.72,10.16). There was no significant effect of order for neither patient-rated CARE (F(1.34,18,76)=1.07, P=0.34) ηp2=0.07) nor clinician-rated CARE (F(1.82,25.42), P=0.63, ηp2=0.03). b) An ANCOVA indicated that higher therapeutic alliance (CARE scores) at intake positively predicted evaluations of the relationship ('HRS score') at the subsequent Clinical-Interaction MRI, across a range of items related to the relationship and social interaction. This association was evident for both for the patient-rated (F(1,144)=9.33, P=0.003 ηp2=0.06) and the clinician-rated (F(1,160)=12.24, P<0.001, ηp2=0.07) scales, indicating the relationship and rapport established at the intake was successfully carried over to the Clinical-Interaction MRI, which was completed on a separate day. There were no main effects or statistical interactions involving 'HRS Item' for neither patients nor clinicians (P's>0.39), suggesting the association between therapeutic alliance at intake Vs relationship at MRI, as well as HRS scores overall, was not different depending on HRS items. Error bars represent Standard error of the mean; [a]Reversed score; CARE=Consultation And Relational Empathy scale.

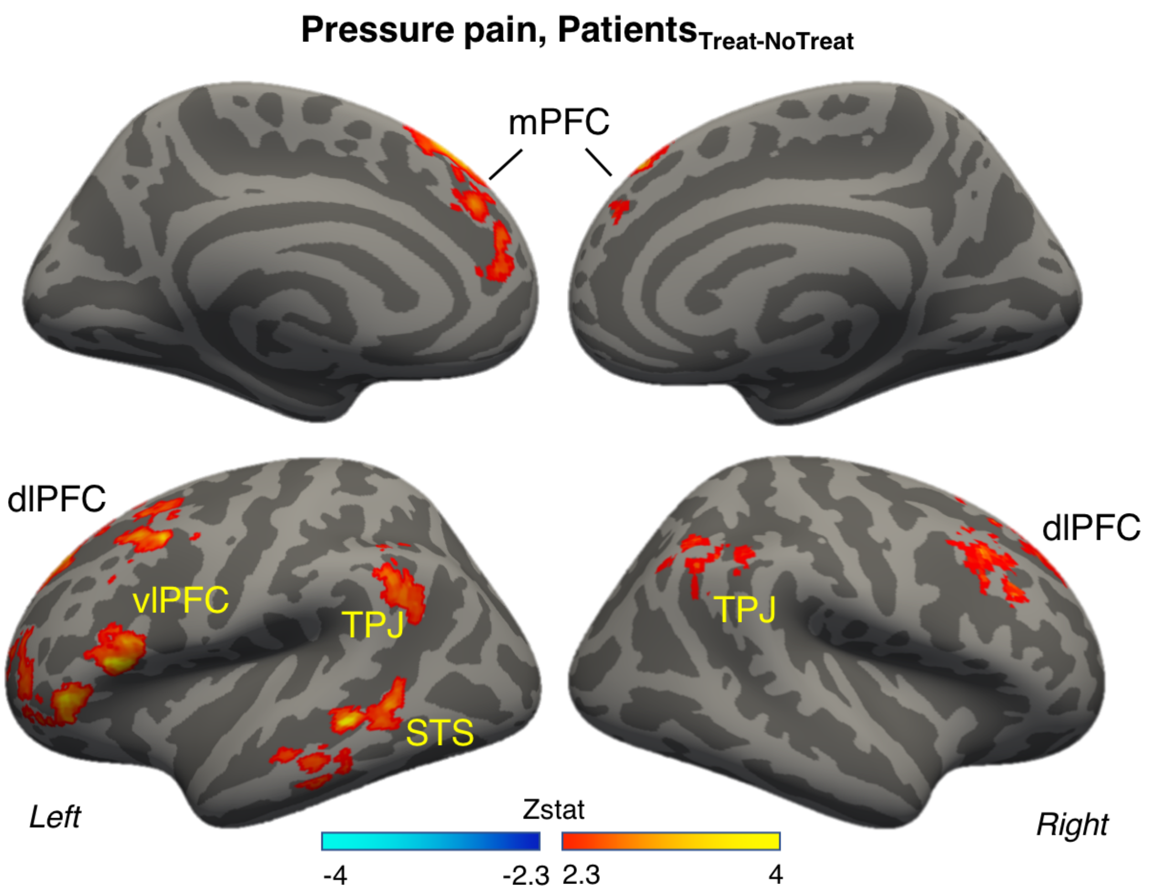


**Fig. S2 | Group map displaying areas where patients showed significant treatment-related (Treat – NoTreat) increase in brain response to pressure pain, thresholded at Z=2.3, P=0.05, cluster-corrected for multiple comparisons.** A whole-brain GLM demonstrated increased fMRI activation of bilateral vlPFC, TPJ, dlPFC, and mPFC, in addition to left STS for treated, relative to nontreated, pain. There were no significant effects in the opposite direction (NoTreat – Treat). This is consistent with a recent meta-analysis of experimental fMRI studies investigating placebo analgesia (*17*), showing that while placebo analgesia was seen in all studies, there was only a marginal BOLD reduction of pain-processing circuitry (*18*). mPFC=medial Prefrontal Cortex; dlPFC=dorsolateral Prefrontal Cortex; vlPFC=ventrolateral Prefrontal Cortex; TPJ=Temporoparietal Junction; STS=Superior Parietal Junction.

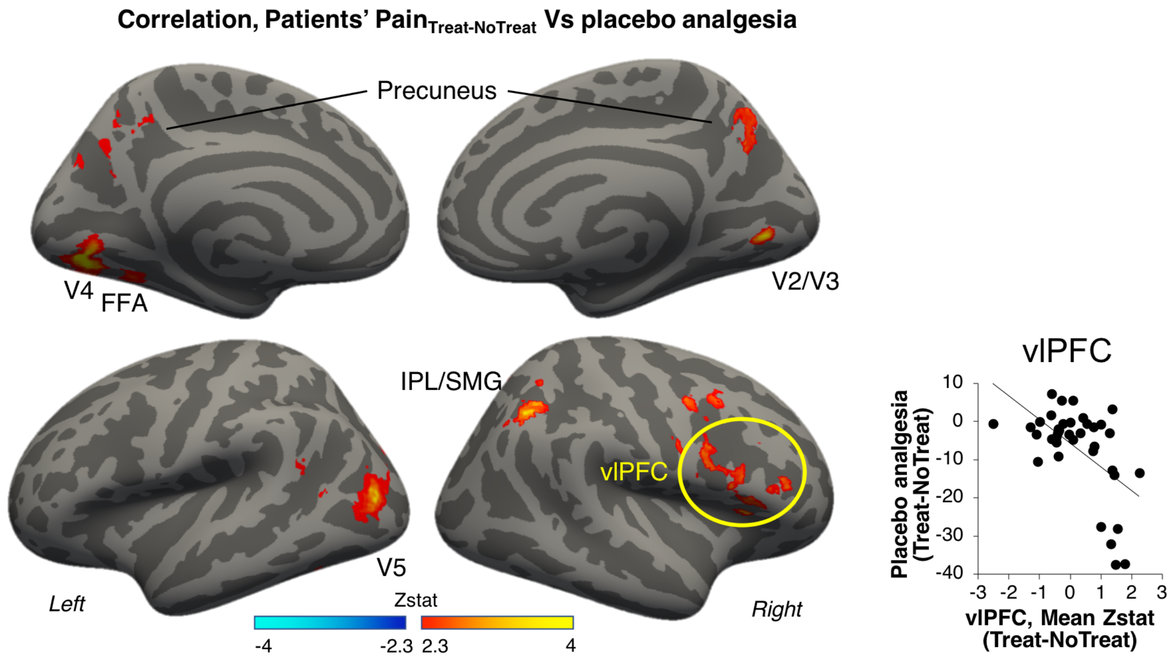


**Fig. S3 | Group map showing areas where stronger treatment-related increases (Treat-NoTreat) in patients' brain response to pressure pain was significantly associated with stronger analgesia (NoTreat-Treat, pain ratings), as indicated by a whole-brain regression analysis.** The displayed group map was thresholded at Z=2.3, P=0.05, cluster-corrected for multiple comparisons. V2-5=Visual areas 1-5; FFA=Fusiform Face Area; IPL=Inferior Parietal Lobule; SMG=Supramarginal Gyrus; vlPFC=ventrolateral Prefrontal Cortex.

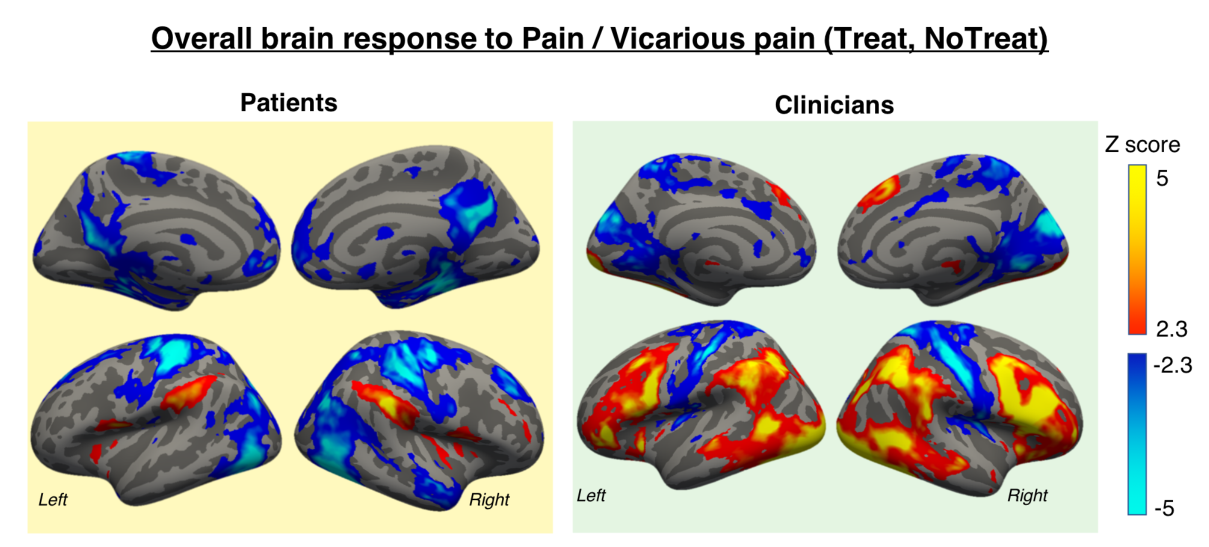


**Fig. S4 | Group maps showing overall response to pain (patients, left) and vicarious pain (clinicians, right), collapsed over Treat/NoTreat conditions.** The displayed group maps were thresholded at Z=2.3, P=0.05, cluster-corrected for multiple comparisons.

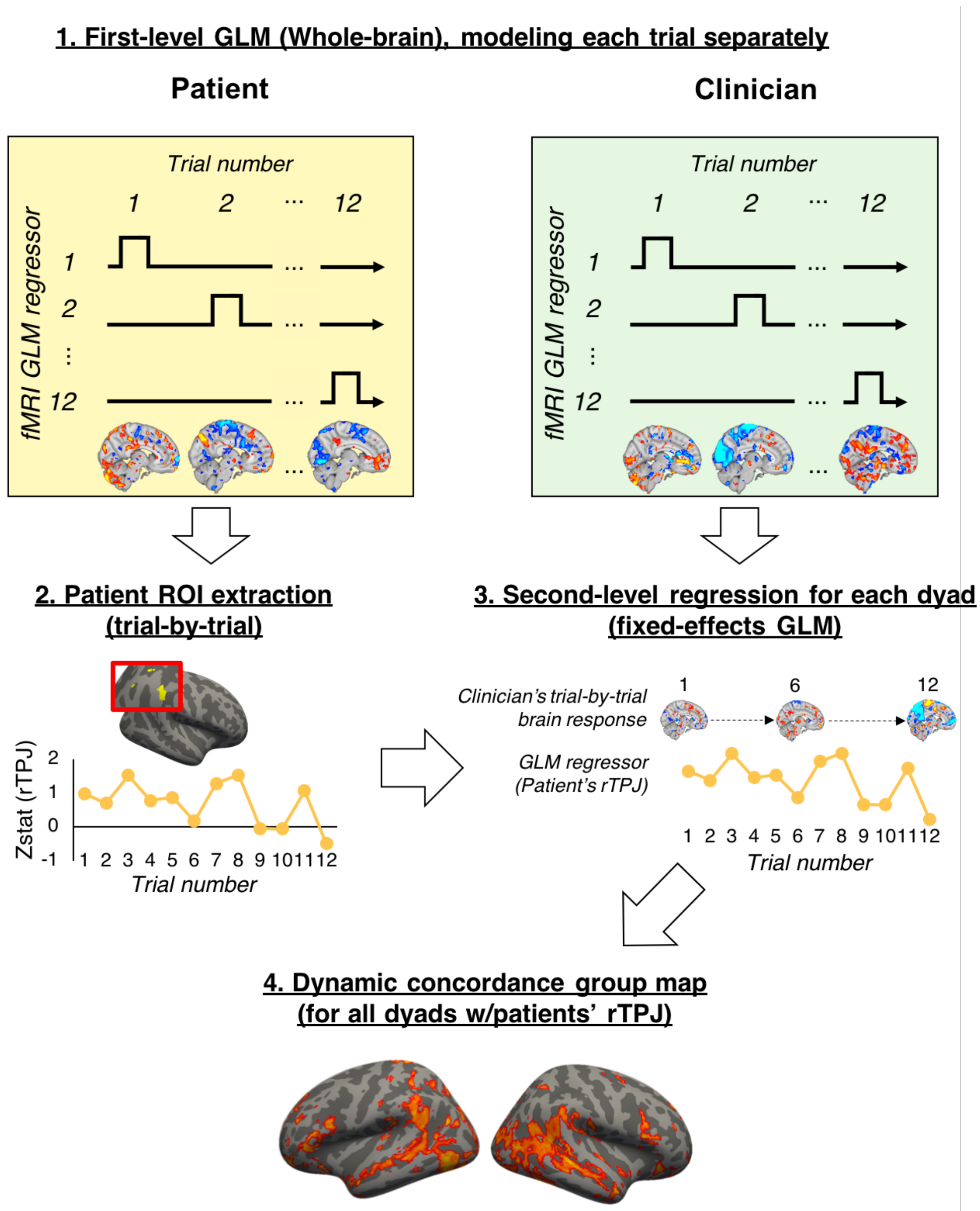


**Fig. S5 | Dynamic concordance analysis pipeline.** 1. To investigate dynamic (time-varying) concordance between patients and clinicians, we first performed a first-level GLM for each subject, where each trial (anticipation phase) was modeled as a separate regressor. This resulted in 12 voxel-wise whole-brain maps (one per trial). 2. For each trial, mean Zstat values were extracted from the patient's social mirroring ROIs (e.g. rTPJ), as identified by a patient-clinician conjunction analysis (see Fig. a). 3. These ROI scores were then used as a regressor in a second-level fixed-effects GLM for the clinician's trial-by-trial brain response during anticipation, which produced a whole-brain map of regions where the clinician's brain response showed dynamic (time-varying) concordance with the patient's social mirroring circuitry (e.g. rTPJ). 4. These steps were taken for each individual dyad, and dynamic concordance maps across dyads were then combined for group analyses. ROI=Region of Interest; rTPJ= right Temporoparietal Junction.

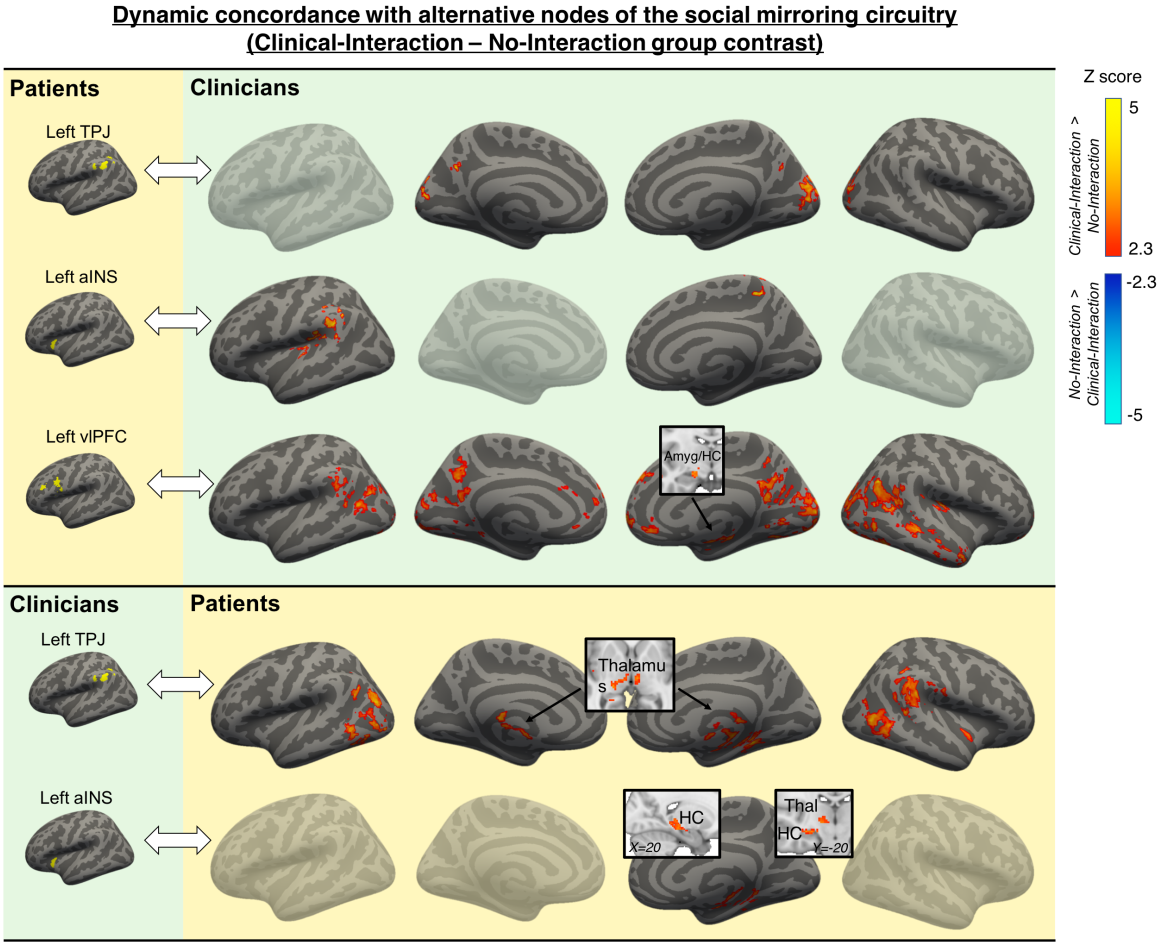


**Fig. S6 | Dynamic concordance with alternative nodes of the social mirroring circuitry.** Each row shows a group contrast (Clinical-Interaction – No-Interaction) of dynamic concordance with different social mirroring ROIs from the partner, as identified by the patient-clinician conjunction group analysis. The top 3 rows show clinician whole-brain concordance with patients' social mirroring ROIs. Patients' left TPJ showed increased concordance with clinicians' precuneus and medial visual cortex for Clinical-Interaction relative to No-Interaction. For the same contrast, patients' left aINS showed increased concordance with clinicians' left TPJ, left posterior insula, and right S1. Left vlPFC showed increased concordance with clinicians' left TPJ, right amygdala/hippocampus, bilateral STS, right mid/posterior insula, vmPFC/pregenual ACC (pgACC), dmPFC, lateral/medial visual cortex, and precuneus. The bottom 2 rows show patient whole-brain concordance with clinicians' social mirroring ROIs. Clinicians' left TPJ showed increased concordance with patients' right TPJ, right aINS, right STS, lateral visual cortex, and bilateral thalamus for Clinical-Interaction relative to No-Interaction. Left aINS showed increased concordance with the right HC and the right thalamus. None of these contrasts showed significant differences in the opposite direction (No-Interaction – Clinical-Interaction). No other nodes of the social mirroring circuitry showed differences in concordance between Clinical-Interaction and No-Interaction. The displayed group maps were thresholded at Z=2.3, P=0.05, cluster-corrected for multiple comparisons. There were no significant effects in the opposite direction (No-Interaction – Clinical-Interaction). TPJ=Temporoparietal Junction; aINS=anterior Insula; vlPFC=ventrolateral Prefrontal Cortex; Amyg=Amygdala; HC=Hippocampus.
